# Fibroblast signaling influences macrophage-dependent, biomaterial-induced tissue remodeling

**DOI:** 10.64898/2026.04.24.720640

**Authors:** Jessica L. Stelzel, Stuart J. Bauer, Bobby Y.X. Ni, Zhi-Cheng Yao, Victor M. Quiroz, Jamie L. Hernandez, Bryan L. McCarty, Russell A. Martin, Kailei D. Goodier, Valerie W. Wong, Sashank K. Reddy, Hai-Quan Mao, Joshua C. Doloff

## Abstract

The ability to induce tissue regeneration on demand using biomaterials remains a major goal in biomedical research, yet significant challenges persist. Among the most advanced biomaterial models, the nanofiber-hydrogel composite has demonstrated a striking ability to induce soft adipose tissue remodeling at the injection site without incorporating exogenous biological cues.^1,2^ However, the underlying mechanisms that drive such a tissue response remain unclear. Here, we show that biomaterial-induced tissue remodeling is driven by sustained and controlled inflammation mediated by macrophages in strong communication with fibroblasts. Notably, both pro-inflammatory and anti-inflammatory signals remained elevated during this process in the long-term, challenging the prevailing notion that inflammation opposes remodeling. Using macrophage depletion in mice, we demonstrate that macrophages are essential for this process. Single-cell RNA sequencing further revealed robust fibroblast-to-macrophage signaling, contrasting with the conventional macrophage-to-fibroblast paradigm, and identified unique *Spp1*⁺ macrophages and *Ctla2a*⁺ fibroblasts within the remodeling niche. These findings provide a comprehensive view of the immune landscape in biomaterial-induced tissue remodeling, highlighting key cellular interactions, prolonged kinetics, and unexpected signaling pathways. By defining key targets and fundamental principles, this work has broad implications for advancing biomaterial-induced tissue regeneration.

## Introduction

The goal of tissue engineering is to regenerate and restore lost or damaged tissue without inducing fibrotic scarring. Except for the liver, mammals have a limited capacity to regenerate or regrow lost tissue. The term tissue remodeling encompasses both the formation of neo-tissue and the reorganization of existing tissues, which can be either non-pathological or pathological. Non-pathological examples include bone remodeling due to exercise,^3^ uterine remodeling during menstruation and pregnancy,^4^ and adipose remodeling in response to nutritional changes.^5^ Pathological examples include airway remodeling in asthma,^6^ vascular remodeling in hypertension,^7^ and cardiac remodeling after myocardial infarction.^8^ Scarring is a fibrotic form of tissue remodeling that aids in wound repair and restores some function, but it is ultimately deleterious as it does not fully restore the tissue to its original state. This process involves the deposition of fibrous connective tissue and is primarily mediated by fibroblasts.

The introduction of biomaterials into the body typically triggers a foreign body response (FBR), which ultimately leads to fibrotic encapsulation, thereby isolating the biomaterials and impairing their functionality. This is especially problematic for biomedical devices that require direct interaction with host tissues, such as implantable sensors, tissue scaffolds, and drug delivery systems. Classical FBR begins with the adsorption of proteins onto the biomaterial surface and the formation of a provisional matrix.^9^ This is followed by inflammation, recruitment of innate immune cells such as macrophages, foreign body giant cell formation, and the deposition of collagen by activated fibroblasts.^9^

Macrophages responding to biomaterials are involved in a variety of host responses, including inflammation and its regulation, phagocytosis, the formation of foreign body giant cells, and the facilitation of tissue remodeling and repair.^10^ Research on the immune response to biomaterials has predominantly emphasized macrophages, whereas other cellular contributors remain comparatively understudied. Although fibroblasts are classically known as matrix-producing effectors of wound healing, fibrosis, and foreign body encapsulation, they are increasingly recognized as active participants in immune regulation. Fibroblasts can sense danger signals, secrete immunomodulatory mediators, shape leukocyte recruitment, and establish stromal niches that influence immune cell function.^11, 12^ However, their immune contributions to biomaterial-induced host response remain incompletely understood. Although crosstalk between macrophages and fibroblasts is increasingly appreciated in this context,^13, 14^ biomaterials studies have often emphasized how macrophages regulate fibroblasts,^15–17^ leaving the reciprocal influence of fibroblasts on macrophages and the broader immune response less well defined.

To counteract the undesirable consequences of fibrotic encapsulation, immune-evasive biomaterials have been designed to mitigate the FBR. Strategies for immune evasion include the modulation of surface topography,^18^ highly adhesive surfaces,^19^ altering size and shape,^20^ as well as the controlled release of immunosuppressive agents.^21^ In contrast, the approach of engineering biomaterials to actively engage with and program the immune system has emerged. While the immune response has traditionally been viewed cautiously due to its association with negative outcomes such as chronic inflammation and fibrosis, recognition of its nuanced role in tissue repair and immunoregulation is growing. Granular biomaterials, like the one studied herein, have been employed to promote tissue integration and repair through their architecture^2, 22^ and, more recently, to actively engage the immune system.^1, 23, 24^

Recently, we developed a nanofiber-hydrogel composite (NHC) with the striking ability to induce remodeling of the injected area into soft adipose tissue without the addition of exogenous biological or biochemical cues.^1, 2^ This NHC consists of a hyaluronic acid hydrogel covalently crosslinked to electrospun poly(ε-caprolactone) nanofiber fragments. This adipogenic remodeling phenomenon is unique among biomaterials and holds significant promise for tissue engineering applications. NHC materials have previously been shown to induce regenerative outcomes in multiple species and injury models.^2, 25, 26^ Additionally, this material has been proven safe in humans in a Phase 1 clinical trial conducted in Europe (NCT04839484). This study investigates the programming of host response toward pro-remodeling and regenerative outcomes to elucidate the mechanisms involved in biomaterial-induced tissue remodeling. Using NHC injected into C57BL/6 mice, we examined the contributions of innate and adaptive immunity to biomaterial-induced tissue remodeling. To causally implicate specific cell types, we employed targeted immune perturbations via multiple genetic knockout and targeted depletion models with a C57BL/6 background. To further identify enriched pathways and infer cellular communication networks, single-cell RNA sequencing was conducted on tissues injected with either the remodeling NHC or the non-remodeling hyaluronic acid hydrogel control.

## Results

### Evaluating biomaterial-induced tissue remodeling: angiogenesis, adipogenesis, & collagen deposition

NHC materials have previously been shown to induce remodeling of the field of injected biomaterial into soft adipose tissue.^1, 2^ To understand the mechanism by which this occurs, NHC materials and hyaluronic acid hydrogel controls were synthesized, mechanically fragmented, and then subcutaneously injected *in vivo* into C57BL/6 mice (**Fig. 1a**). This immunocompetent mouse strain was selected because of its intact fibrotic response that mimics that of humans.^18, 27^ For initial characterization experiments, NHC & gel control materials of different, matched stiffnesses (soft: 150 Pa or stiff: 450 Pa) were tested to show that results are notably more dependent on material type rather than stiffness (**Supplementary Fig. 1**). Vascular branching was easily visible in NHC by day 7, whereas gel controls lacked this feature (**Fig. 1b**). This was assessed with quantitative polymerase chain reaction (qPCR), which showed that NHC materials induced expression of pro-angiogenic factor *Vegfa* (**Fig. 1c**). As this was also observed in the gel control, the clear difference in blood vessel formation may be attributed to the expression differences in the Vegf receptors or to the dilution of signal in the NHC due to immune cell infiltration. In addition to blood vessel formation, the overall remodeling progression of the injected volume into adipose tissue was seen and quantified with Masson’s Trichrome-stained histological cross sections of explanted materials (**Fig. 1d–f, Supplementary Fig. 2**). Adipose tissue formation was first noticeable by 3 months and was robust at 1 year. This transition from sterile biomaterial to adipose tissue was preceded by the formation of a unique “pre-remodeling zone” that appeared by day 35 in NHC materials (**Fig. 1g**). This pre-remodeling zone is an area of dense cellular infiltration characterized by a unique pattern of collagen deposition in a fine scaffold-like network. In Masson’s Trichrome staining, it appears pink/purple with dark purple nuclei and thin blue lines of deposited collagen (**Fig. 1e**). While the FBR typically causes biomaterials to develop a thick outer collagen capsule,^9^ the NHC materials had thin collagen tendrils deposited throughout their volume as they were remodeled (**Fig. 1e, h**). This biomaterial-induced remodeling deviated from the canonical type 3 to type 1 collagen transition typically associated with progressing from early to late wound healing (**Fig. 1i**). In a systematic analysis of all known mouse collagen isoforms, collagen deposition in NHC was marked by higher expression of collagen genes such as *Col6a3*, *Col12a1*, and *Col24a1* (**Fig. 1j, k, Supplementary Fig. 3**). Similar biomaterial-induced tissue remodeling resulting in long-term adipogenesis was observed in another model organism when NHC materials were subcutaneously injected in New Zealand White rabbits (**Supplementary Fig. 4**). Lumina, a commercial-grade NHC material, underwent a remodeling progression similar to what was observed in mice. This progression consisted of significant cellular infiltration followed by remodeling of the NHC into adipose tissue. Radiesse, the commercially available dermal filler used as a control, did not undergo comparable remodeling.

**Figure 1.**
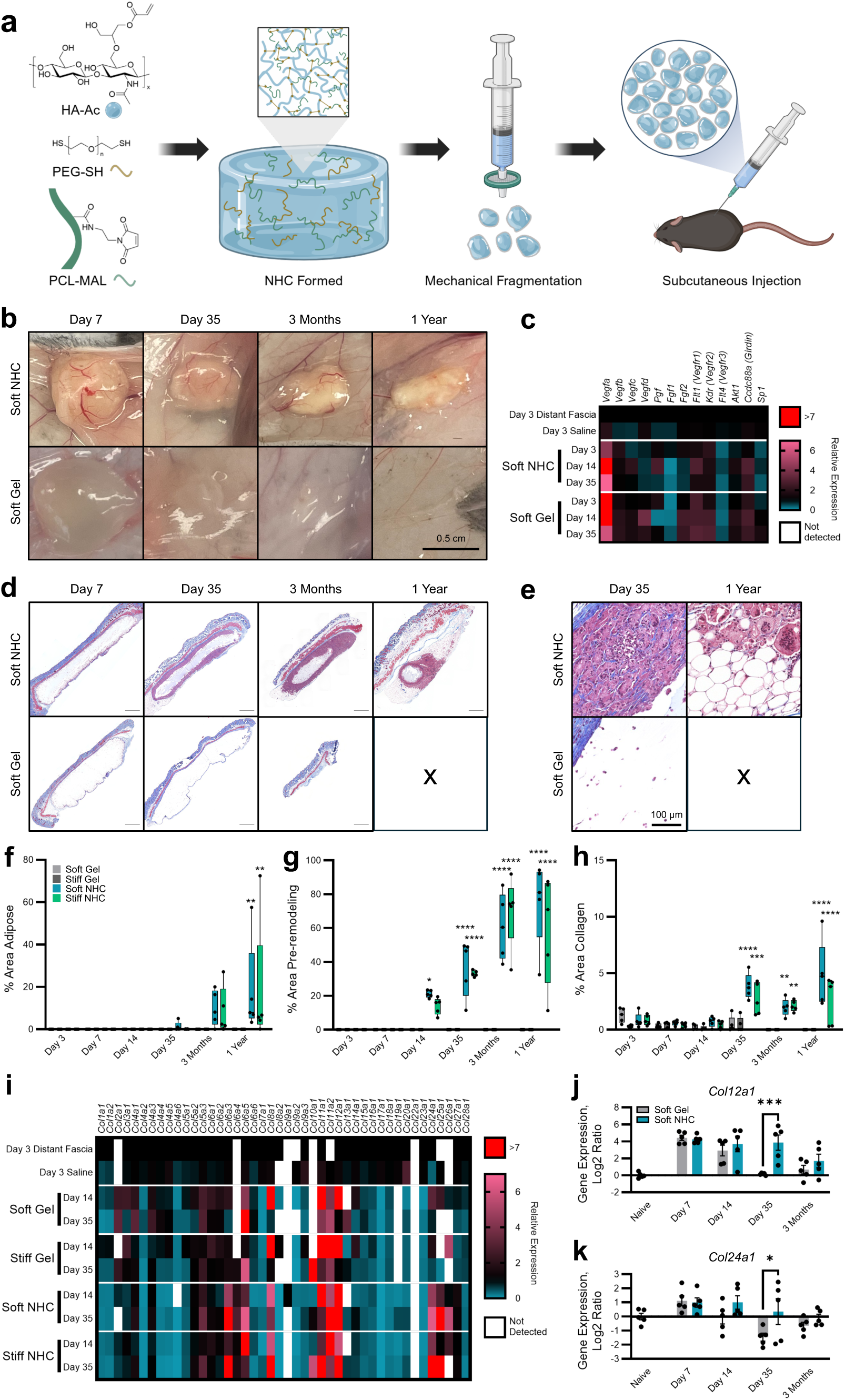
NHC materials as a model of tissue remodeling. **a,** Schematic showing chemical structure (*left*), NHC material preparation (*middle*), and implantation workflow (*right*). NHC components include acrylate-modified hyaluronic acid (HA-Ac), dithiol poly(ethylene glycol) (PEG-SH), and maleimide-functionalized poly(ε-caprolactone) (PCL-MAL) nanofibers. Created using Biorender.com. **b,** Representative photographs of stiffness-matched soft (150 Pa) hyaluronic acid hydrogel (“gel”) and NHC materials about to be explanted from C57BL/6 mice after various timepoints. **c,** Heatmap of qPCR analysis of angiogenesis-related gene expression in explanted materials (soft: 150 Pa) relative to distant fascia in C57BL/6 mice with pooled n = 5 biological replicates. **d, e,** Representative Masson’s Trichrome-stained histologic cross-sections of materials subcutaneously injected into C57BL/6 mice, showing gross morphology including skin (**d**), or zoomed-in view inside materials (**e**). Scale bars are 1000 µm (**d**) or as annotated (**e**). **f–h,** Analysis of Masson’s Trichrome histological images quantifying the amount of adipogenesis (**f**), pre-remodeling zone (**g**), and collagen deposition (**h**) inside the materials at various timepoints. Plots show the percent area of the component of interest from the total area of the biomaterial. Data, min to max box and whisker (n = 5 biological replicates). Two-way ANOVA followed by Tukey’s multiple comparisons test. **i,** Heatmap of qPCR analysis of collagen-related gene expression in explanted materials relative to distant fascia in C57BL/6 mice with pooled n = 5 biological replicates. **j, k,** NanoString gene expression for *Col12a1* (**j**) and *Col24a1* (**k**) on materials explanted from C57BL/6 mice. Plots show the absolute transcript counts normalized to naïve fascia (scale: log2 ratio). Data, mean ± SEM (n = 5 biological replicates). Two-way ANOVA followed by Šídák’s multiple comparisons test. In this figure’s graphs, statistics shown are (*) timepoint and stiffness-matched comparisons of NHC vs. gel control. \**p* < 0.05, \*\**p* < 0.01, \*\*\**p* < 0.001, \*\*\*\**p* < 0.0001.

### Programmed inflammation of recruited immune cells for tissue remodeling

Following characterization, the immunological mechanisms underlying this unique remodeling response were further interrogated. The NHC material recruited significantly more cells than the gel control, as determined through quantification of hematoxylin and eosin (H&E) stained slides (**Fig. 2a–c, Supplementary Fig. 5**). Multi-color flow cytometry determined that for both materials the infiltrating cells largely consisted of macrophages, dendritic cells, neutrophils, natural killer cells, and T cells (**Fig. 2d, Supplementary Figs. 6 and 7, Supplementary Table 1**). The majority of responding CD45^+^ cells were innate rather than adaptive immune cells, which follows logically, as remodeling is related to innate immune system-dominated processes, such as wound healing.^28^ The lack of substantial differences in cellular makeup between the NHC and gel control led to a deeper investigation of transcriptional profiles and functional responses. Through profiling immune-related gene expression, an unexpected pattern of sustained and controlled immune activation was observed in the NHC. Interestingly, while expression of many immune-related genes peaked between days 7 and 35, the NHC induced higher expression of both predominantly pro-inflammatory (e.g., *Il1b*,^29^ *Il1a*,^29^ *Tnf*^30^) and anti-inflammatory (e.g., *Arg1*,^31^ *Tgfb1*,^32^ *Il10ra*^33^) genes out to long-term timepoints (**Fig. 2e, Supplementary Fig. 8**). Along with the lack of skin discomfort or irritation in the mice, this implies continued immune activation but not deleterious chronic inflammation. This is best visualized at the 3-month timepoint, where the NHC group showed ongoing elevated expression of immune-related genes, while the gel control returned to baseline. Representative genes *Il1a*, *Tgfb1*, and *Ptprc* demonstrate this significant sustained elevation in the NHC (**Fig 2f–h**). *Il1a* is a cytokine that acts as a key danger signal to initiate inflammation^29^ and *Tgfb1* is a potent immunosuppressive cytokine that contributes to the resolution of inflammation,^32^ demonstrating the ongoing balance. The persistently elevated expression of *Ptprc* (*Cd45*), a pan-immune cell marker, indicates prolonged immune cell infiltration into the NHC material. These long-lasting differences indicate the NHC is actively interfacing with and influencing the immune system as it undergoes remodeling. Additional genes of interest highly expressed by the NHC are *Spp1* (**Fig. 2i**) and *Cxcr4* (**Fig. 2j**). *Spp1* had the highest average expression in the NHC, with very clear and sustained separation between the NHC and gel control by day 7. *Spp1* is associated with increased immune cell infiltration,^34^ driving collagen deposition,^35, 36^ and the modulation of macrophage phenotype.^37^ *Cxcr4* encodes a chemokine receptor with reported roles in cell migration, development, tissue regeneration, proliferation, hematopoiesis, and angiogenesis.^38^

**Figure 2.**
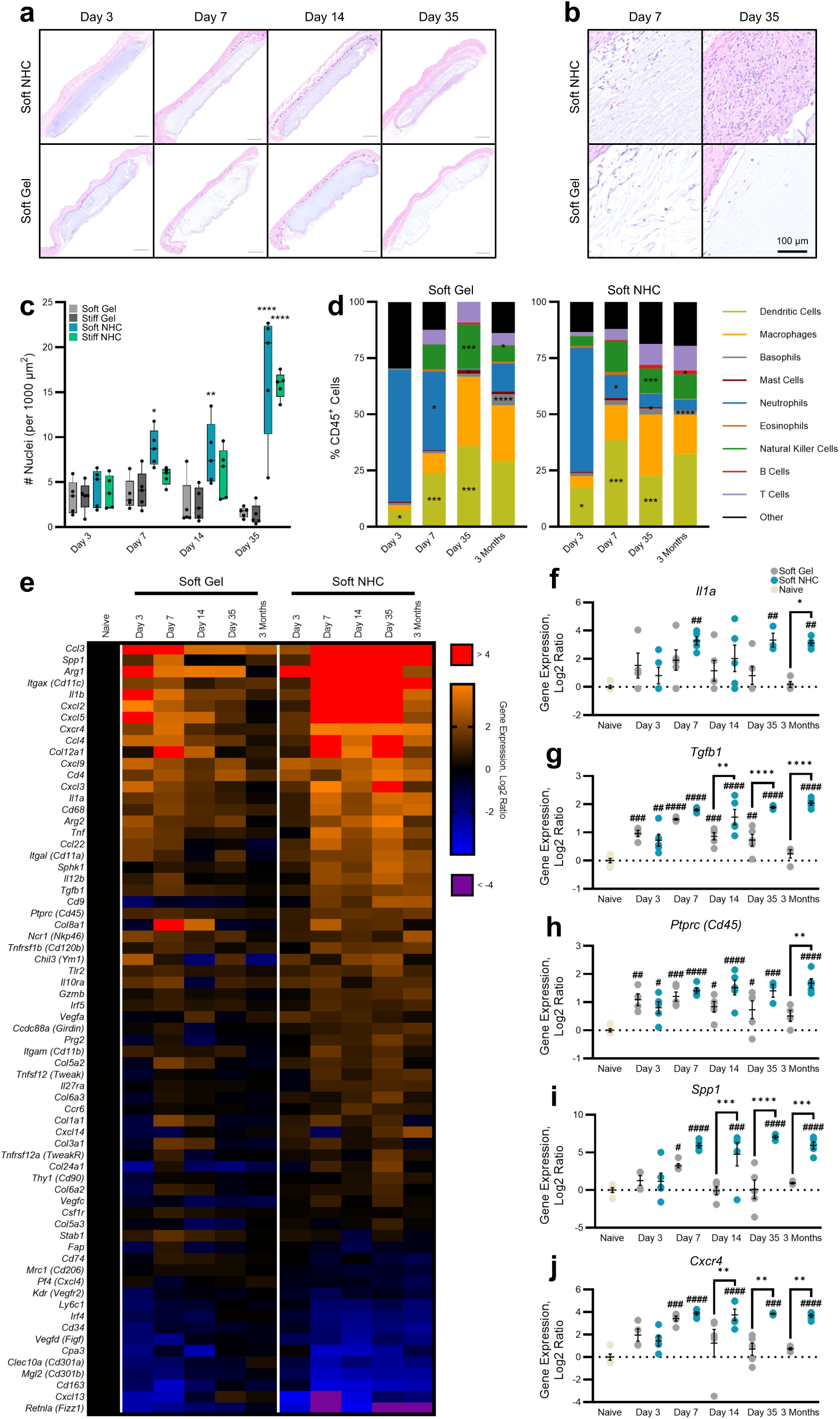
Biomaterial-induced tissue remodeling occurs through sustained, controlled inflammation. **a, b,** Representative H&E-stained histologic cross-sections of stiffness-matched soft (150 Pa) hyaluronic acid hydrogel (“gel”) and NHC materials subcutaneously injected into C57BL/6 mice (**a**), and zoomed-in view inside materials (**b**). Scale bars are 1000 µm (**a**) or as annotated (**b**). These H&E histological images correspond to the Masson’s Trichrome-stained samples in Figure 1. **c,** Analysis of H&E histological images quantifying the amount of cellular infiltration inside the materials (soft: 150 Pa or stiff: 450 Pa) at various timepoints. Plot shows the nuclei per area. Data, min to max box and whisker (n = 5 biological replicates). Two-way ANOVA followed by Tukey’s multiple comparisons test. **d,** Flow cytometric analysis of cells recruited to materials explanted from C57BL/6 mice at various timepoints, shown as %CD45^+^ cells. Data, mean (n = 5 biological replicates). Two-way ANOVA followed by Tukey’s multiple comparisons test. Immune cell types were defined as follows: dendritic cells (CD45^+^CD11c^+^), macrophages (CD45^+^CD11b^+^CD68^+^F4/80^+^), basophils (CD45^+^CD11b^+^FceR1^+^), mast cells (CD45^+^CD11b^-^FceR1^+^), neutrophils (CD45^+^CD11b^+^Ly6g^+^), eosinophils (CD45^+^Cd11b^+^SiglecF^+^F4/80^+^), natural killer cells (CD45^+^NKp46^+^), B cells (CD45^+^CD19^+^CD3^-^), & T cells (CD45^+^CD3^+^CD19^-^). **e,** Heatmap showing NanoString gene expression of significantly differentially expressed genes (adjusted *p*-value < 0.05) for materials explanted from C57BL/6 mice at various timepoints. The heatmap shows absolute transcript counts normalized to naïve fascia (scale: log2 ratio). Data, mean (n = 3-5 biological replicates). All genes shown were significant in a timepoint-matched NHC vs. gel control comparison, with statistics run by nSolver. Genes were ordered based on average expression across all NHC groups. **f–j,** NanoString gene expression for *Il1a* (**f**), *Tgfb1* (**g**), *Ptprc* (**h**), *Spp1* (**i**), and *Cxcr4* (**j**). Plots show the absolute transcript counts normalized to naïve fascia (scale: log2 ratio). Data, mean ± SEM (n = 5 biological replicates). Two-way ANOVA followed by the Holm-Šídák multiple comparisons test. In this figure’s graphs, statistics shown are (*) timepoint and stiffness-matched comparisons of NHC vs. gel control, as well as (^#^) comparisons vs. naïve fascia. *,^#^:*p* < 0.05; **,^##^:*p* < 0.01; ***,^###^:*p* < 0.001; ****,^####^:*p* < 0.0001.

### Macrophages play a causal role in biomaterial-induced tissue remodeling

Cells infiltrating the NHC material were assessed to determine their causal and mechanistic roles in driving tissue remodeling. By using knockout and depletion mouse models to perturb responding immune cells, cell types that altered the progression of tissue remodeling were established. This was quantitatively assessed by comparing the percent pre-remodeling zone of Masson’s Trichome-stained NHC specimens at day 35 between wild-type and knockout/depletion mice (**Fig. 3a, b, Supplementary Fig. 9a**). Importantly, depleting macrophages prevented the formation of the pre-remodeling zone, while depleting neutrophils sped up its formation (**Fig. 3b**). These experiments confirmed that macrophages are necessary for biomaterial-induced tissue remodeling, and interestingly, neutrophil presence slows that remodeling.

**Figure 3.**
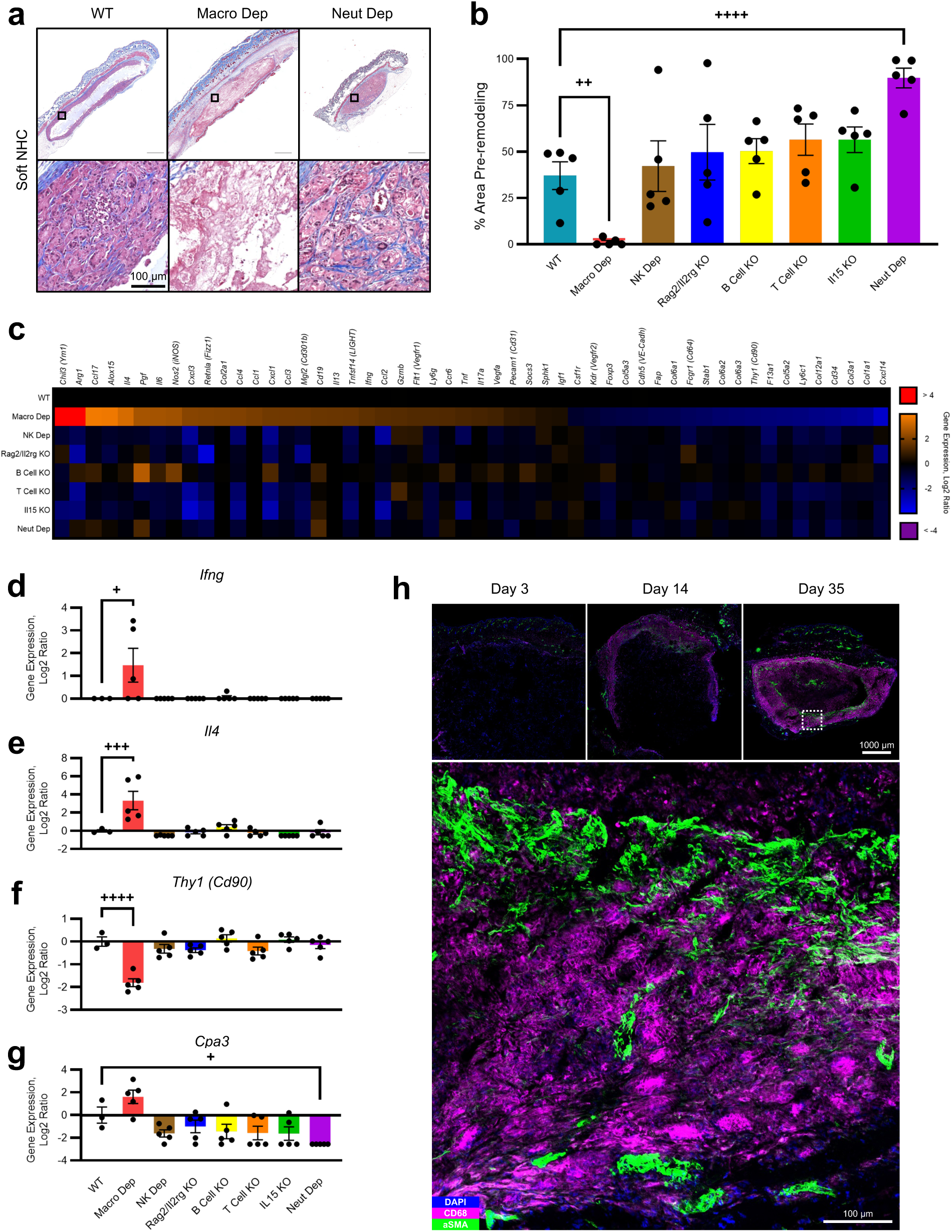
Macrophages causally implicated in biomaterial-induced tissue remodeling. **a,** Representative Masson’s Trichrome-stained histologic cross-sections of soft (150 Pa) NHC materials subcutaneously injected into knockout (KO)/depletion (Dep) mouse models with a C57BL/6 background, in comparison to a fully immune-competent wild-type (WT) control. Materials were explanted at day 35. A full, zoomed-out view (*top row*) and a smaller, zoomed-in view of the matrices (*bottom row*) are shown. Mouse models shown are WT C57BL/6 mice, macrophage depletion (Macro Dep) via Clodrosome injection, and neutrophil depletion (Neut Dep) via anti-Ly6g antibody injection. Scale bars are 1000 µm (*top*) or as annotated (*bottom*). **b,** Analysis of Masson’s Trichrome histological images quantifying the amount of pre-remodeling zone inside explanted NHC materials at d35 in knockout/depletion mouse models. Plot shows the percent area of the pre-remodeling zone from the total area of the biomaterial. Additional mouse models shown are natural killer (NK) cell depletion via anti-asialo GM1 antibody injection into C57BL/6 mice, Rag2/Il2rg double knockout mice, B cell knockout via muMt^-^ mice, T cell knockout via B6-nude mice, and additional NK/T cell dysfunction with Il15 KO mice. Data, mean ± SEM (n = 5 biological replicates). One-way ANOVA followed by Šídák’s multiple comparisons test. **c,** Heatmap showing NanoString gene expression of significantly differentially expressed genes (*p*-value < 0.05) for materials explanted from WT and knockout/depletion mice at day 35. The heatmap shows absolute transcript counts normalized to naïve fascia (scale: log2 ratio). Data, mean (n = 3-5 biological replicates). All genes shown were significant in Macro Dep vs. WT and Macro Dep vs. 2+ other knockout/depletion models, with statistics run by nSolver. Genes were ordered based on expression in the Macro Dep group. **d–g,** NanoString gene expression for *Ifng* (**d**), *Il4* (**e**), *Thy1* (**f**), and *Cpa3* (**g**). Plots show the absolute transcript counts normalized to naïve fascia (scale: log2 ratio). Data, mean ± SEM (n = 3-5 biological replicates). One-way ANOVA followed by the Holm-Šídák multiple comparisons test. **h,** Representative immunofluorescence images of histologic cross-sections of NHC materials subcutaneously injected into C57BL/6 mice and explanted at various timepoints. Staining was performed using DAPI, CD68, and αSMA. Scale bars are as annotated. In this figure’s graphs, statistics shown are (^+^) comparisons of knockout/depletion mice vs. WT mice. ^+^*p* < 0.05; ^++^*p* < 0.01; ^+++^*p* < 0.001; ^++++^*p* < 0.0001.

When surveying the immune landscape, significant increases in the expression of predominantly pro-inflammatory genes (e.g., *Tnf*,^30^ *Ifng*,^39^ and various chemokines) and predominantly anti-inflammatory genes (e.g., *Il4*^40^ and *Il13*^40^) were observed in the non-remodeling macrophage depletion model compared to wild type (**Fig. 3c–e, Supplementary Fig. 9b**). This suggests that the NHC’s macrophages were involved in immunoregulation. Notably, *Il4* and *Il13* are both considered signature type 2 cytokines.^40^ Their increase may be a repair-associated response to clodrosome-induced cell death which was rendered functionally nonproductive due to the absence of macrophages. This macrophage depletion-induced shift in the immune system likely disrupted the delicate balance necessary for tissue remodeling. The macrophage depletion model also had decreased expression of collagen and fibroblast-related genes (*Thy1*, *Cd34*, and multiple collagen isoforms), likely due to the collagen-laden pre-remodeling zone not forming (**Fig. 3c, f**). Neutrophil depletion significantly decreased the mast cell marker *Cpa3*, likely due to neutrophil-driven mast cell recruitment, whereas it was elevated in the macrophage depletion model (**Fig. 3g**). The opposing trends in macrophage and neutrophil depletion suggest a minor role for these rare cells.

Macrophages, which were causally indicated in biomaterial-induced tissue remodeling, were observed infiltrating the NHC via CD68 immunofluorescence staining (**Fig. 3h**). Their infiltration over time roughly corresponded with the development of the pre-remodeling zone. Immunofluorescence staining for alpha-smooth muscle actin (αSMA), a fibroblast marker, showed strong activity in the NHC material that is particularly notable along the inner edge of the macrophage-laden pre-remodeling zone (**Fig. 3h**).

### Enriched immune and remodeling pathways revealed alongside a fibroblast-dominated signaling network

To investigate remodeling at a higher resolution, single-cell RNA sequencing (scRNA-seq) was performed on NHC and gel control materials retrieved at days 7 and 35 post-injection. Sequencing resulted in 52,329 high-quality cells, from 3 to 4 mice in each group, at an average depth of 79,620 reads per cell. Following standard preprocessing steps, unsupervised clustering identified 11 distinct clusters. Cycling cells and doublets were excluded, leaving 9 clusters. Some of the identified clusters of interest were further divided into subclusters, and additional doublets were removed. Overall, this clustering provided higher resolution of the previously identified populations, as well as revealed additional infiltrating cell types of fibroblasts, monocytes, and innate lymphoid cells (**Fig. 4a, Supplementary Fig. 10a–c**). Neutrophils were only detected in the NHC via scRNA-seq, despite being present in both materials as determined by flow cytometry (**Fig. 2d, Supplementary Fig. 10c, d**). Neutrophil scRNA-seq analysis is challenging due to their inherently low RNA content and sensitivity to degradation.^41^ Additionally, neutrophil differences between groups, such as variations in transcriptional activity or maturation, could have contributed to their degradation and therefore lack of detection in the gel control.^41^ To identify transcriptomic differences across samples, differential gene expression (DGE) analysis was performed. The comparisons between the remodeling NHC and the gel control at both timepoints had a large number of significant differentially expressed genes (DEGs) (6,302 genes at day 7; 8,205 genes at day 35), indicating very different biological processes had occurred (**Supplementary Fig. 10e, f**). This demonstrates a stark difference between the NHC undergoing tissue remodeling and the mild FBR response elicited by the gel control.

**Figure 4.**
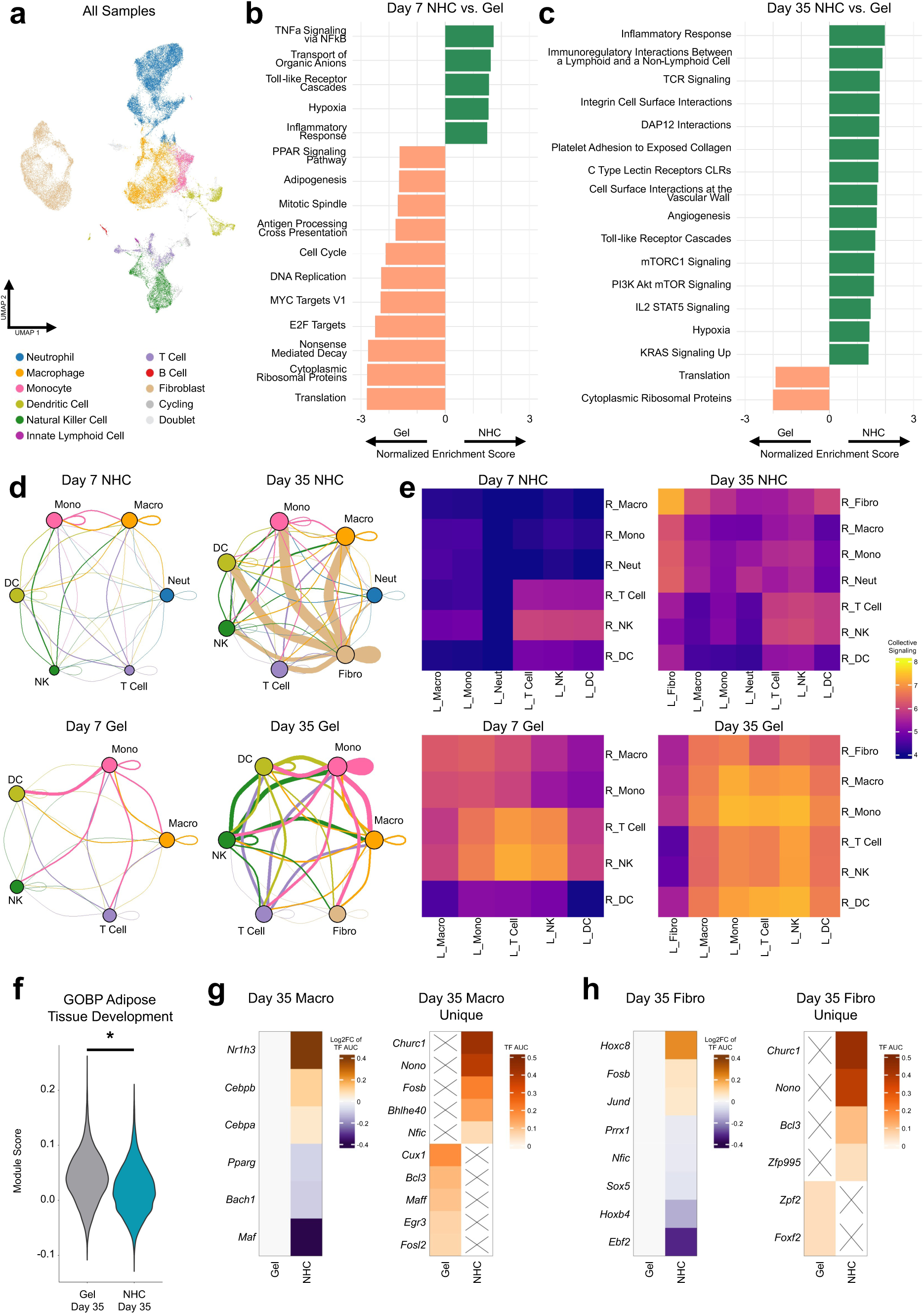
scRNA-seq uncovered remodeling-related pathways and fibroblast-centric inferred signaling network. **a,** UMAP plot showing annotated scRNA-seq clusters for all samples sequenced, 52,329 cells total. Samples are soft (150 Pa) NHC and hyaluronic acid hydrogel (“gel”) control materials, subcutaneously injected into C57BL/6 mice and explanted at days 7 and 35 (n = 3 or 4 biological replicates per group). **b, c,** Normalized enrichment scores of select significantly enriched gene sets (adjusted *p*-value < 0.05) from Hallmark, BioCarta, WikiPathways, and Reactome collections. NHC vs. gel control comparisons based on pseudobulk differential gene expression are for day 7 (**b**) and day 35 (**c**). **d,** Visual representation of inferred signaling networks via the dominoSignal package for all conditions. Nodes represent clusters, and node size corresponds to the total incoming signaling to each cluster. Edges illustrate signaling from one cluster to another. Edge color is based on the cluster expressing the ligands, and edge thickness corresponds to outgoing signaling (summed ligand expression) from the node of the same color to the other node it connects to. **e,** Heatmaps of inferred outgoing signaling from sending clusters (denoted “L_” for ligands) to receiving clusters (denoted “R_” for receptors) for all conditions. Color is based on the collective (summed) ligand signaling score of the sending cluster. **f,** Violin plot of module scores by material type at day 35 for the “GOBP Adipose Tissue Development” pathway. Independent t-test. **g, h,** Heatmaps of cluster-averaged pySCENIC-calculated transcription factor area under the curve (TF AUC) scores for selected adipogenesis-related transcription factors found by DominoSignal in both the NHC and gel control (scale: log2 fold change) (*left*) or in only one material group (*right*). Transcription factors shown were activated in macrophages by fibroblasts (**g**) or activated in fibroblasts by macrophages (**h**). In this figure’s graphs, statistics shown are (*) timepoint and stiffness-matched comparisons of NHC vs. gel control. \**p* < 0.05, \*\**p* < 0.01, \*\*\**p* < 0.001, \*\*\*\**p* < 0.0001.

To determine pathways implicated in remodeling, ranked gene set enrichment analysis (GSEA) was performed (**Fig. 4b, c**). Significantly enriched gene sets for the gel controls overall mainly involved protein translation and the cell cycle. The NHC’s overall significantly enriched gene sets are related to immune activation, immunoregulation, angiogenesis, collagen, and signaling. Pathways heavily influenced by the lack of neutrophils in the gel control were excluded (**Supplementary Fig. 10g**). Importantly, a balance of different types of immune activation is demonstrated by the significant immune-related NHC gene sets, such as toll-like receptor cascades, TNFα signaling via NF-κB, and immunoregulatory interactions between a lymphoid and a non-lymphoid cell (**Fig. 4b, c**). These findings reinforce that NHC-induced remodeling occurs through an immune response that is sustained but still controlled. Other significant signaling pathways included mTORC1, PI3K/Akt/mTOR, and KRAS, which showcase an increased capacity for proliferation, growth, and survival in the infiltrating cells remodeling the NHC (**Fig. 4b, c**). Angiogenesis, a hallmark of the NHC and required process for remodeling, was also reflected in the NHC’s GSEA results. Hypoxia was significant in the NHC by day 7, and other vasculature-related gene sets, such as cell surface interactions at the vascular wall and angiogenesis, became significant at day 35 (**Fig. 4b, c**). Interestingly, TCR signaling and IL2 STAT5 signaling were both significant in the NHC at day 35, indicating T cell activity (**Fig. 4b, c**). However, the T cell knockout suggested their involvement did not play a causal role (**Fig. 3b**).

Given the multitude of interesting cell types and the complexity of the response, an investigation into cell communication was undertaken by inferring receptor-ligand-transcription factor networks from the scRNA-seq data. This analysis revealed distinct inferred signaling networks in both the NHC and gel control (**Fig. 4d, e, Supplementary Fig. 11**). Interestingly, while the gel control did not result in remodeling, at both timepoints it had more outgoing signaling (by number of activated signaling pathways and average ligand expression). However, with that said, the comparatively lower outgoing NHC signaling was surprisingly dominated by fibroblasts signaling to themselves, as well as to neutrophils, monocytes, and macrophages. Of note, macrophages and neutrophils were found to play important roles in the progression of remodeling via murine depletion models (**Fig. 3b**). Despite having a larger proportion of fibroblasts, the gel control had very little inferred signaling originating from fibroblasts (**Fig. 4e**). This stark difference in fibroblast signaling, paired with the fact that the NHC fibroblasts communicated primarily with implicated immune cell types, suggests an active immunological role for fibroblasts in tissue remodeling. This immune role of fibroblasts is explored further in a later section.

### Early signaling in NHC’s macrophages and fibroblasts tied to the negative regulation of adipogenesis

Counterintuitively, at day 35 multiple adipogenesis-related gene sets were significantly elevated in the gel control, which did not induce adipogenesis, compared to the adipogenic NHC material (**Fig. 4f, Supplementary Fig. 12a**). As substantial adipogenesis in the NHC was not observed until 3 months (**Fig. 1d–f**), activity at the early day 35 timepoint likely reflects the signaling involved in establishing a pro-adipogenic niche rather than adipogenesis itself. Adipogenesis-related pathways were significantly elevated primarily in macrophages and fibroblasts (**Supplementary Fig. 12b–h**), demonstrating their early regulation of the NHC’s long-term remodeling outcome. When signaling between the macrophages and fibroblasts was examined, many of the top transcription factors were adipogenesis-related (**Fig. 4g, h, Supplementary Fig. 13a, b**). Interestingly, the adipogenesis-related transcription factors higher in NHC macrophages and fibroblasts were predominantly related to the negative regulation of adipogenesis. For example, the NHC’s macrophages and fibroblasts both had increased activation of the *Fosb*, whose splice product ΔFosB is a highly stable transcription factor that prevents adipogenesis and interferes with early adipogenic commitment.^42^ The NHC’s macrophages also had increased activation of *Bhlhe40* and *Nr1h3*, transcription factors which can inhibit adipogenic differentiation.^43, 44^ When examining the gene sets from the implicated anti-adipogenic and adipogenesis-regulating transcription factors, the pattern of being significantly elevated in the NHC is evident in both the macrophages and fibroblasts (**Supplementary Fig. 13c, d**). Notably, TGF-β superfamily members *Inhba* and its receptors *Acvr1b* and *Acvr2a* were all significantly increased in the NHC’s macrophages, with *Acvr1b* also significantly increased in the NHC’s fibroblasts. This signaling functions as a brake on early adipogenesis, keeping adipocyte progenitors more proliferative and less differentiated.^45^ Additional notable genes significantly higher in the NHC include *Runx2*, a negative regulator of adipogenic commitment,^46^ *Pten*, which reduces the proliferation and differentiation of adipogenic progenitors,^47^ and a variety of Wnt pathway genes. This early regulatory signaling activity may represent a preparatory phase that permits the establishment of a permissive pro-adipogenic niche before adipogenesis begins. Niche formation is multi-faceted: around day 35 in NHC the pre-remodeling zone was being established (**Fig. 1g**), a fine network of collagen was being deposited throughout its volume (**Fig. 1e, h, Supplementary Fig. 13e**), numerous immune cells were infiltrating (**Fig. 2c, d**), and signaling to negatively regulate adipogenesis was occurring (**Fig. 4f–h, Supplementary Fig. 12a, Supplementary Fig. 13a–d**). Overall, the early signaling in the NHC that eventually leads to adipogenesis was interestingly anti-adipogenic.

### Macrophage subclustering highlights *Ms4a7*^+^ and *Spp1*^+^ Macrophages

Special attention was paid to macrophages due to the depletion data demonstrating their causal role in the observed NHC-induced tissue remodeling (**Fig. 3b**). When performing unsupervised subclustering of the scRNA-seq data, 5 distinct macrophage/monocyte subclusters emerged: *Ms4a7*^+^ macrophages, *Spp1*^+^ macrophages, *Retnla*^+^ macrophages, low-quality macrophages, and monocytes (**Fig. 5a, b, Supplementary** Fig. 14). Due to quality control metrics, one macrophage subcluster was excluded due to its high percentage of mitochondrial reads, which are associated with cell damage and stress (**Supplementary** Fig. 15a, b). When splitting the cells by sample group, the relative abundance of the subclusters varied heavily (**Fig. 5c**). Differential abundance analysis showed that the *Spp1*^+^ macrophages were essentially only present in the NHC, while the *Retnla*^+^ macrophages were far more abundant in the gel control (**Fig. 5d, e**). Recall, *Spp1* was the gene with the highest average expression in the NHC out to 3 months in the NanoString data, underscoring its importance in this system (**Fig. 2e**). Through examining marker genes and performing trajectory analysis, the NHC was found to contain more monocytes and monocyte-derived-like macrophages, most notably *Spp1*^+^ macrophages, while the gel control contained more tissue resident-like macrophages (**Supplementary Fig. 16a–i**). This data corresponds with the increased immune infiltration observed in the NHC materials (**Fig. 2c**). Additionally, the macrophage subclusters do not conform to the M1/M2 macrophage polarization binary, with each subcluster expressing at least one classic M1-like gene and one classic M2-like gene (**Fig. 5f**).

**Figure 5.**
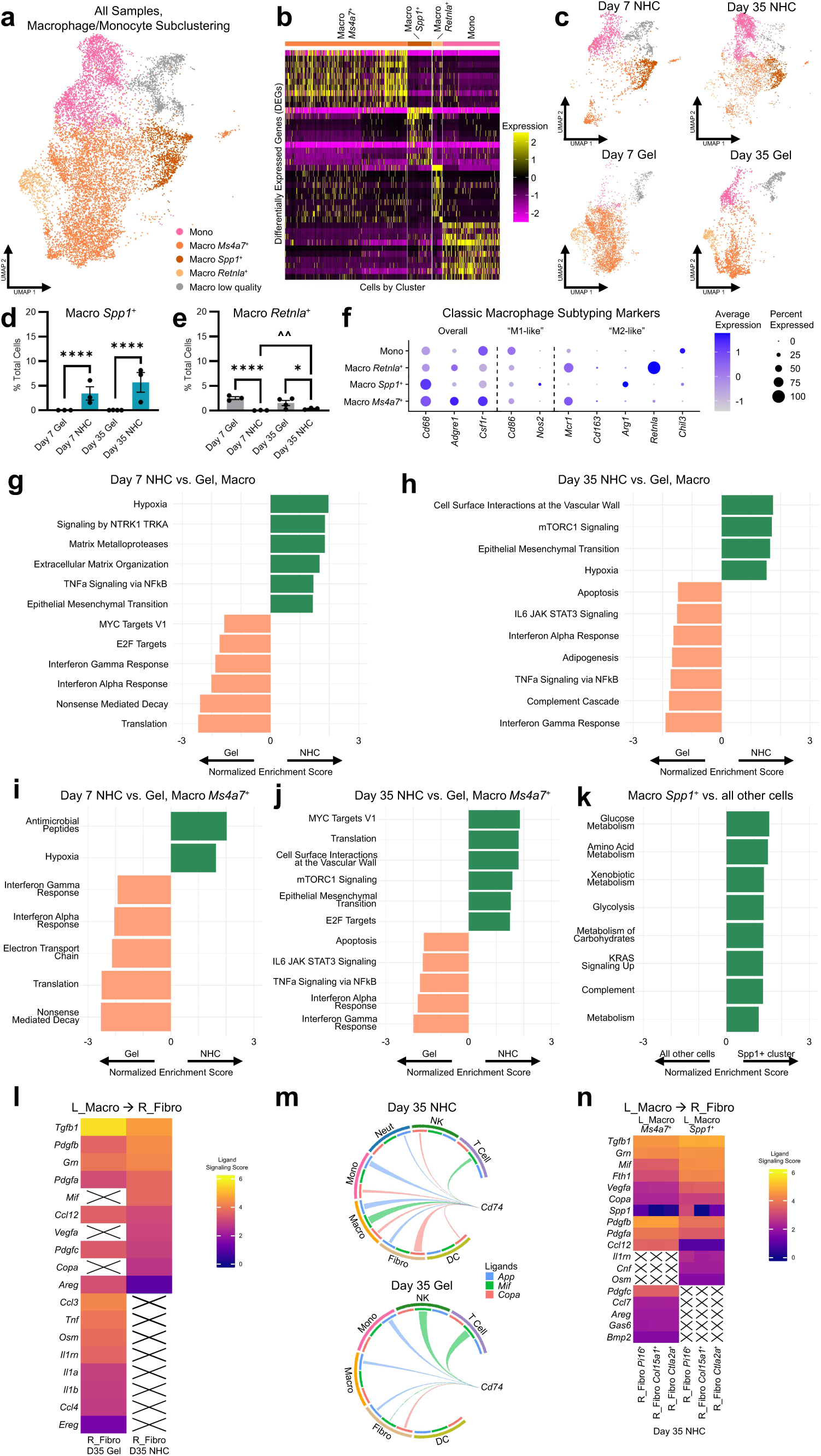
Macrophage subclustering and signaling showcases remodeling-associated subclusters and signaling to fibroblasts. **a,** UMAP plot showing macrophage and monocyte subclustering across all conditions, 12,277 cells total. **b,** Heatmap of significant differentially expressed genes (DEGs) with the highest log fold change per subcluster. **c,** UMAP plots of cell subtypes recruited to materials split by condition (timepoint + material) for macrophages and monocytes. **d, e,** Differential abundance results comparing cluster abundance between groups for *Spp1*^+^ macrophages (**d**) and *Retnla*^+^ macrophages (**e**), found using edgeR’s generalized linear model framework. Data, mean ± SEM (n = 3 or 4 biological replicates). **f,** Dot plot depicting normalized expression of classic macrophage subtyping marker genes for macrophage/monocyte subclusters. **g–k,** Normalized enrichment scores of select significantly enriched gene sets (adjusted *p*-value < 0.05) from Hallmark, BioCarta, WikiPathways, and Reactome collections. NHC vs. gel control comparisons based on pseudobulk differential gene expression are for day 7 macrophages (**g**), day 35 macrophages (**h**), day 7 *Ms4a7*^+^ macrophages (**i**), and day 35 *Ms4a7*^+^ macrophages (**j**). Cluster vs. all other cells comparison based on the Wilcoxon rank-sum test differential gene expression is for *Spp1*^+^ macrophages (**k**). **l,** Heatmap of inferred outgoing signaling from the macrophage cluster to the fibroblast cluster at day 35 for NHC and gel control, showing selected ligands with the largest differences in ligand signaling scores. Sending cluster(s) are denoted “L_” for ligands and receiving cluster(s) are denoted “R_” for receptors. Color is based on ligand signaling scores. An X represents the lack of a ligand-receptor linkage. **m,** Chord plot showing expression of ligands that can activate a specific receptor, *Cd74*. Chord width represents mean ligand expression in the originating cluster. Day 35 plots shown for NHC (*top*) and gel control (*bottom*). **n,** Heatmap of inferred outgoing signaling from the macrophage subclusters to the fibroblast subclusters at day 35 for NHC, showing selected ligands with the largest differences in ligand signaling scores. Sending cluster(s) are denoted “L_” for ligands and receiving cluster(s) are denoted “R_” for receptors. Color is based on ligand signaling scores. An X represents the lack of a ligand-receptor linkage. In this figure’s graphs, statistics shown are (*) timepoint and stiffness-matched comparisons of NHC vs. gel control, as well as (^^^) comparisons between the same group at different timepoints. *,^^^:*p* < 0.05; **,^^^^:*p* < 0.01; ***,^^^^^:*p* < 0.001; ****,^^^^^^:*p* < 0.0001.

DGE was performed on the macrophage cluster and its subclusters, revealing numerous significant gene expression differences between the NHC and gel control macrophages (2,497 genes at day 7; 3,530 genes at day 35) (**Supplementary Fig. 17a–e**). The DGE results were then used to identify significantly enriched pathways through GSEA. NHC macrophages, determined to be causal in biomaterial-induced tissue remodeling, were involved in angiogenesis, epithelial-to-mesenchymal transition (EMT), extracellular matrix (ECM) organization, and induced inflammation via TNFα/NF-κB signaling, as well as stimulated proliferation via mTORC1 and TrkA signaling (**Fig. 5g, h**). Of note, the gene sets for hypoxia and EMT were significant in NHC macrophages at both timepoints. While macrophages themselves cannot undergo EMT as they are not epithelial cells, they can drive the process. Multiple significant gene sets in the NHC macrophages were also significant in the NHC overall, indicating that the overall NHC enrichment of hypoxia, mTORC1 signaling, and cell surface interactions at the vascular wall was presumably contributed by the macrophages. In contrast, gel macrophages were involved in translation, interferon responses, proliferation, and complement (**Fig. 5g, h**). The macrophage subcluster GSEA results highlighted their distinctions (**Fig. 5i–k**). The NHC’s *Ms4a7*^+^ macrophage cluster’s significant gene sets denoted association with angiogenesis, proliferation, mTORC1 signaling, and EMT (**Fig. 5i, j**). In contrast, the gel *Ms4a7*^+^ macrophages were identified as having roles in interferon responses, protein translation, and signaling (**Fig. 5i, j**). *Spp1*^+^ macrophages, which were only substantially present in the NHC samples, were highly metabolically active as well as involved in KRAS signaling and complement (**Fig. 5k**). *Spp1*^+^ macrophages are an interesting subtype implicated in fibrosis,^35, 36^ wound healing,^48^ and cancer.^37, 49^ *Retnla*^+^ macrophages, which were mostly present in the gel control, did not have any significantly enriched gene sets.

### NHC-induced macrophage signaling to fibroblasts involves angiogenesis, immunomodulation, and migration

Since fibroblasts were a prominent component of the NHC signaling network and macrophages were causally implicated in biomaterial-induced tissue remodeling, macrophage signaling to fibroblasts was further investigated (**Fig. 5l–n, Supplementary Fig. 17f–k**). Because there were not enough fibroblasts present for analysis by day 7, fibroblast-related analysis was performed at day 35 only. The macrophages were signaling more in the gel control, both overall and specifically to fibroblasts, but they only caused tissue remodeling in the NHC (**Fig. 4e**). The ligands with the largest differences in NHC vs. gel control macrophage-to-fibroblast signaling revealed differences in the skew of the immune response and angiogenesis (**Fig. 5l**). In the gel control, macrophages signaling to fibroblasts had higher ligand expression levels of predominantly pro-inflammatory cytokines (*Il1a*,^29^ *Il1b*,^29^ *Tnf*^30^) and chemokines (*Ccl3*, *Ccl4*, *Ccl12*),^50^ further highlighting the macrophage differences between the NHC and gel control. *Tgfb1*, which is known to promote fibrotic fibroblast phenotypes,^32, 51^ was also substantially increased in the gel macrophage signaling (**Fig. 5l**). Conversely, the NHC macrophages had higher ligand expression levels of *Pdgfa*, *Pdgfb*, and *Vegfa*, growth factors that play a significant role in blood vessel development,^52^ as well as immune-related ligands *Grn*, *Mif*, and *Copa* (**Fig. 5l**). All six of these macrophage-expressed ligands primarily targeted fibroblasts (**Supplementary Fig. 17f–g**). *Grn* is implicated in immunoregulation, proliferation, and wound repair^53^; *Mif* is a cytokine associated with increased inflammation, proliferation, and tissue regeneration^54^; and *Copa* is involved in intracellular protein transport, whereas mutations in this gene cause immune dysregulation.^55^ Lastly, both *Mif* and *Copa* are ligands for *Cd74*, which had more signaling in day 35 NHC than the gel (**Fig. 5m, Supplementary Fig. 17h**). Mif-Cd74 is a high-affinity interaction that promotes immune activation and tissue repair across various organs.^56^

At the subcluster level, *Ms4a7*^+^ and *Spp1*^+^ macrophages had similar overall signaling patterns in NHC samples, but showed distinct differences when zooming into specific ligands. Key ligands distinguishing the subclusters are *Mif*,^54, 56^ *Fth1*,^57^ and *Tgfb1*^32, 51^ in NHC’s *Spp1*^+^ macrophages, suggesting a regenerative, immunomodulatory, and fibrotic role; and *Pdgfa*,^52, 58, 59^ *Pdgfb*,^52, 58, 59^ *Pdgfc*,^52, 58, 59^ and *Ccl12*^60^ in the NHC *Ms4a7*^+^ macrophages, suggesting functions in angiogenesis, wound healing, fibrosis, and immune cell recruitment (**Fig. 5n, Supplementary Fig. 17i–k**). This suggests that both *Ms4a7*^+^ and *Spp1*^+^ macrophages played a role in the unique remodeling response induced by the NHC. Notably, *Spp1* ligand expression varied depending on the targeted fibroblast subcluster, with most *Spp1*-*Cd44* signaling occurring between *Spp1*^+^ macrophages and *Pi16*^+^ fibroblasts (**Fig. 5n**). *Spp1* signaling can enhance fibroblast differentiation,^61^ proliferation,^62^ and migration^62^ as well as drive EMT^63^—key processes for tissue remodeling. This is particularly relevant because Pi16 marks progenitor fibroblasts.^64^ Additionally, *Spp1*^+^ macrophages have been shown to influence fibroblasts by promoting a more fibrotic phenotype.^36^

### Stark differences in fibroblast abundance and subtypes between NHC and control

Fibroblasts are of interest due to their well-established role in wound healing, the unique collagen deposition observed in remodeling NHC materials, and their robust signaling patterns in this model. Fibroblast outgoing signaling at day 35 was predominant in NHC and, in contrast, minimal in the gel control. In the day 35 NHC group, fibroblast signaling primarily targeted the NHC’s most important cell types: macrophages, monocytes, neutrophils, and fibroblasts. The fibroblasts were separated into three subclusters: *Pi16*^+^ fibroblasts, *Col15a1*^+^ fibroblasts, and *Ctla2a*^+^ fibroblasts (**Fig. 6a, b, Supplementary Fig. 14**). Although fibroblasts were scarce at day 7, they were clearly abundant by day 35 (**Fig. 6c, d**). At day 35, the gel control had a significantly higher abundance of *Pi16*^+^ fibroblasts, *Col15a1*^+^ fibroblasts, and fibroblasts overall, but the *Ctla2a*^+^ fibroblasts were nearly exclusive to the NHC (**Fig. 6d–g**).

**Figure 6.**
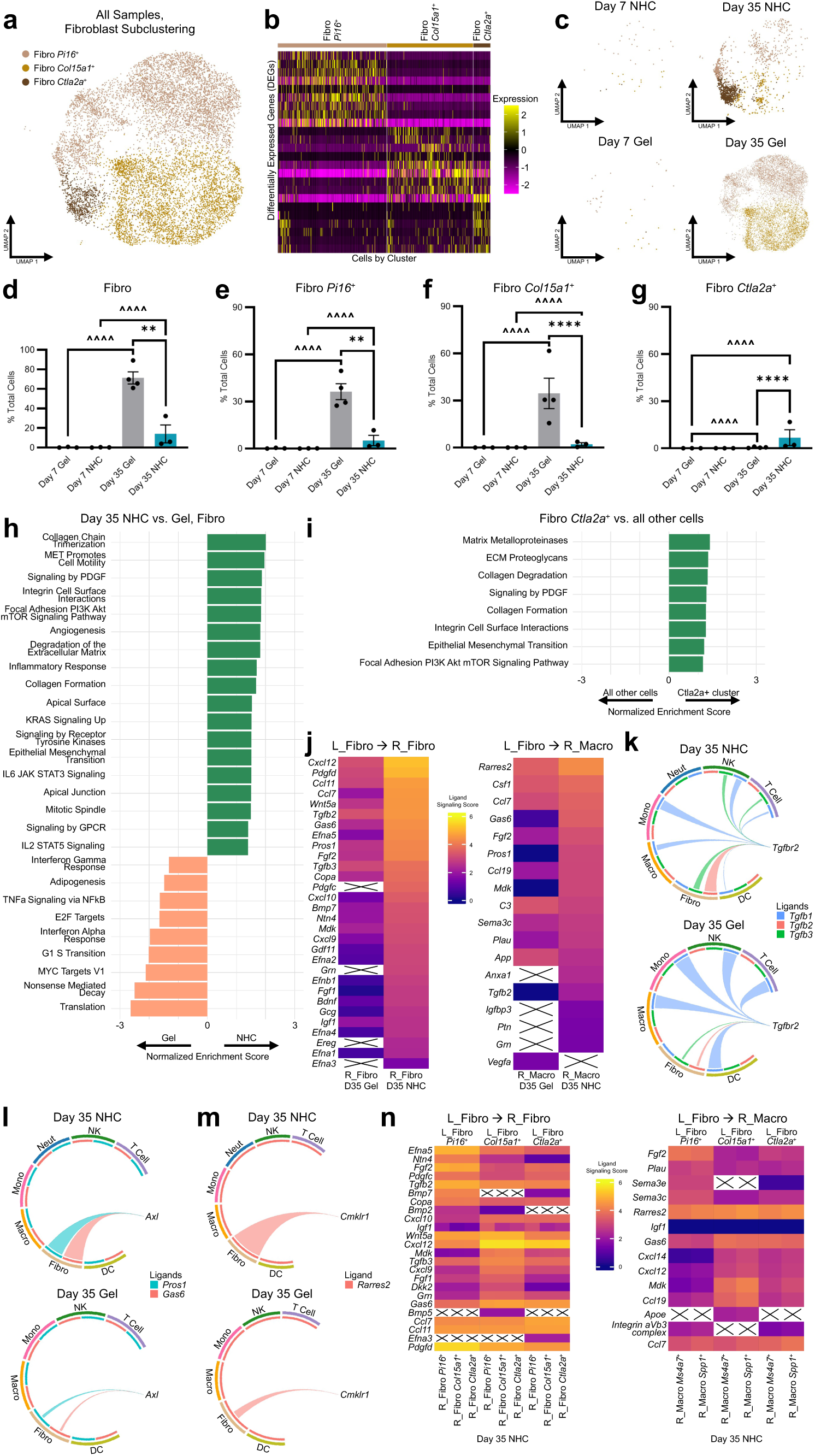
Fibroblast subclustering and signaling reveal them as a bridge between the immune response and tissue remodeling. **a,** UMAP plot showing fibroblast subclustering across all conditions, 10,734 cells total. **b,** Heatmap of significant differentially expressed genes (DEGs) with the highest log fold change per subcluster. **c,** UMAP plots of cell subtypes recruited to materials split by condition (timepoint + material) for fibroblasts. **d–g,** Differential abundance results comparing cluster abundance between groups for all fibroblasts (**d**), *Pi16*^+^ fibroblasts (**e**), *Col15a1*^+^ fibroblasts (**f**), and *Ctla2a*^+^ fibroblasts (**g**), found using edgeR’s generalized linear model framework. Data, mean ± SEM (n = 3 or 4 biological replicates). **h, i,** Normalized enrichment scores of select significantly enriched gene sets (adjusted *p*-value < 0.05) from Hallmark, BioCarta, WikiPathways, and Reactome collections. NHC vs. gel control comparison based on pseudobulk differential gene expression is for day 35 fibroblasts (**h**). Cluster vs. all other cells comparison based on the Wilcoxon rank-sum test differential gene expression is for *Ctla2a*^+^ fibroblasts (**i**). **j,** Heatmaps of inferred outgoing signaling from the fibroblast cluster to the receiving cluster, fibroblasts (*left*) or macrophages (*right*), at day 35 for NHC and gel control, showing selected ligands with the largest differences in ligand signaling scores. Sending cluster(s) are denoted “L” for ligands and receiving cluster(s) are denoted “R_” for receptors. Color is based on ligand signaling scores. An X represents the lack of a ligand-receptor linkage. **k–m,** Chord plots showing expression of ligands that can activate a specific receptor. Chord width represents mean ligand expression in the originating cluster. Day 35 plots shown for NHC (*top*) and gel control (*bottom*) for receptors *Tgfbr2* (**k**), *Axl* (**l**), & *Cmklr1* (**m**). **n,** Heatmaps of inferred outgoing signaling from the fibroblast subclusters to the receiving subclusters, fibroblasts (*left*) or macrophages (*right*), at day 35 for NHC, showing selected ligands with the largest differences in ligand signaling scores. Sending cluster(s) are denoted “L” for ligands and receiving cluster(s) are denoted “R_” for receptors. Color is based on ligand signaling scores. An X represents the lack of a ligand-receptor linkage. In this figure’s graphs, statistics shown are (*) timepoint and stiffness-matched comparisons of NHC vs. gel control, as well as (^^^) comparisons between the same group at different timepoints. *,^^^:*p* < 0.05; **,^^^^:*p* < 0.01; ***,^^^^^:*p* < 0.001; ****,^^^^^^:*p* < 0.0001.

Similarly to the overall and macrophage DGE, the NHC vs. gel control fibroblast DGE also resulted in numerous significant DEGs (1,598 genes at day 35), showcasing the distinctions between the fibroblasts in the NHC and gel control (**Supplementary Fig. 18a**). At day 35, the gel control fibroblasts’ significantly enriched gene sets indicated their involvement in translation, proliferation, and signaling (**Fig. 6h**). Their signaling included elevated expression of gene sets related to interferons and TNFα. Conversely, the NHC fibroblasts had an abundance of significantly enriched gene sets of interest in remodeling. They were involved in ECM degradation, collagen formation, angiogenesis, adhesion, motility, EMT, and multiple signaling pathways (**Fig. 6h**). These enriched pathways reveal the NHC fibroblasts’ role as highly mobile cells that invade and remodel the biomaterial. Some significant gene sets in the NHC fibroblasts were also significant in the NHC overall, such as integrin cell surface interactions, angiogenesis, KRAS signaling up, and IL2 STAT5 signaling (**Fig. 4c & Fig. 6h**). This indicates that these NHC vs. gel control differences were presumably driven by the fibroblasts. Furthermore, the enriched gene sets in the NHC fibroblasts aligned well with those in the NHC macrophages, indicating that they contribute to similar processes and potentially work together; both cell types exhibited significant GSEA results related to angiogenesis, the ECM, and EMT. Angiogenesis and ECM-related gene sets are consistent with the blood vessel formation and unique spatial collagen deposition observed in the remodeling NHC. EMT is a developmental process where epithelial cells transform into migratory mesenchymal cells such as fibroblasts.^65^ It has also been observed in wound healing and cancer,^65^ and now in biomaterial-induced tissue remodeling as well.

Some of the significantly enriched gene sets in the NHC *Pi16*^+^ fibroblasts at day 35 related to collagen, angiogenesis, adhesion, motility, wound healing, and signaling (**Supplementary Fig. 18b, c**). These GSEA results were extremely similar to those from the overall fibroblasts, highlighting the *Pi16*^+^ fibroblast subset as a key driver of the overall NHC vs. gel control differences (**Fig. 6h**). The *Col15a1*^+^ fibroblasts had very few significantly enriched pathways (**Supplementary Fig. 18d, e**). Conversely, the *Ctla2a*^+^ fibroblasts abundant in the NHC had multiple significantly enriched gene sets. These gene sets—which involved ECM remodeling, adhesion, EMT, and signaling—were also similar to those from the overall day 35 NHC fibroblasts, underscoring the importance of the *Ctla2a*^+^ subset (**Fig. 6h, i, Supplementary Fig. 18f**). These *Ctla2a*^+^ fibroblasts are also *Postn*^+^, and previous work has implicated communication between *Postn*^+^ fibroblasts and *Spp1*^+^ macrophages in fibrotic deposition.^36^

### Strong fibroblast signaling tied to remodeling, immunomodulation, and adipogenesis

Fibroblast signaling was the most distinct difference between the NHC and gel control signaling networks. In day 35 NHC, fibroblast outgoing signaling was highest towards themselves, with high levels also directed towards neutrophils, monocytes, and macrophages (**Fig. 4e**). Since fibroblast signaling was surprisingly predominant and macrophages were causally implicated in this biomaterial-induced tissue remodeling, fibroblast-to-fibroblast and fibroblast-to-macrophage signaling were probed in more depth (**Fig. 6j–n, Supplementary Fig. 19a–e**). Although the gel control had a significantly higher abundance of fibroblasts than the NHC, fibroblasts had the highest outgoing signaling in the NHC and lowest outgoing signaling in the gel control (**Fig. 4e, Fig. 6d**). The most distinct ligands involved in fibroblast-fibroblast signaling, defined as having the largest NHC vs. gel control expression differences, were all increased in the NHC (**Fig. 6j**). Similarly, all but 3 of the most distinct ligands in fibroblast-to-macrophage signaling were also elevated in the NHC over the gel control. Among others, the fibroblast-to-fibroblast ligands which exhibited the most pronounced NHC vs. gel control differences included growth factors (*Tgfb2*, *Tgfb3*, *Bmp7*, *Gdf11*, *Fgf2*, *Fgf1*, *Pdgfd*, *Pdgfc*, *Mdk*, *Grn*, & *Igf1*), chemokines (*Cxcl12*, *Ccl11*, *Ccl7*, *Cxcl10*, & *Cxcl9*), and ephrins (*Efna5*, *Efna2*, *Efnb1*, *Efna4*, *Efna1*, & *Efna3*). Notably, ephrins play essential roles in development, migration, and immunity, while also serving as key regulators of tissue repair and fibrosis. For fibroblast-to-macrophage signaling, the ligands with the greatest NHC vs. gel control differences included multiple cytokines (*Csf1*, *Ccl7*, *Ccl19*) and growth factors (*Fgf2*, *Mdk*, *Tgfb2*, *Ptn*, *Grn*). Fibroblast signaling to both fibroblasts and macrophages involved ligands associated with tissue remodeling, angiogenesis, immunomodulation, migration, and adipogenesis. These functions are demonstrated by the *Tgfbr2, Cmklr1,* and *Axl* receptors (**Fig. 6k–m, Supplementary Fig. 19c–e**).

In the gel control, signaling to the *Tgfbr2* receptor was predominantly engaged by *Tgfb1* produced by multiple cell types, with very little *Tgfb2* or *Tgfb3* (**Fig. 6k**). In NHC, the *Tgfbr2* receptor received more communication from *Tgfb2* and *Tgfb3*, which were mainly produced by fibroblasts. While all Tgfb isoforms are involved in fibrosis/wound healing and development and can induce EMT, they have differential effects.^51^ *Tgfb1* and *Tgfb2* are classically considered pro-fibrotic and *Tgfb3* anti-fibrotic, but their influence is more complex and context-dependent.^51^ The NHC’s increased *Tgfb2* and *Tgfb3* represent a nuanced shift in fibro-modulation, potentially underlying the change from the FBR to remodeling. Another ligand of interest is *Gas6*, which was a top NHC-enriched ligand for fibroblast outgoing signaling (**Fig. 6l**). *Gas6* is the primary ligand for the Axl receptor, which drives EMT, is associated with stem cell maintenance, and promotes angiogenesis.^66^ The Axl receptor has also been shown to promote an immunoregulatory microenvironment by influencing the recruitment and polarization of immune cells, including macrophages.^66^ Axl signaling represents an avenue by which the NHC fibroblasts could directly affect macrophage phenotype, therefore influencing remodeling. *Rarres2* encodes the adipocytokine Chemerin, the primary ligand for the Cmklr1 receptor, and was the most expressed ligand in NHC fibroblast-to-macrophage signaling (**Fig. 6m**). Rarres2-Cmklr signaling is implicated in adipogenesis, angiogenesis, promoting repair, and leukocyte recruitment.^67^ Further, NHC fibroblasts exhibited increased expression of ligands that are both drivers and suppressors of adipogenesis (*Rarres2*,^67^ *Bmp7*,^68^ *Igf1*,^69^ *Wnt5a*,^70^ *Tgfb2*,^71^ *Tgfb3*^71^), demonstrating regulation of this long-term remodeling outcome as early as day 35 (**Fig. 6j**).

While differences in fibroblast subcluster abundance were clear, their signaling differences were more subtle and complex. The fibroblast subclusters appeared similar on the overall signaling heatmap, albeit with the *Col15a1*^+^ fibroblasts having slightly stronger signaling to a few additional clusters and the *Ctla2a*^+^ fibroblasts having slightly weaker signaling to other fibroblasts (**Supplementary Fig. 11**). Based on ligand expression differences, the most characteristic ligands for each fibroblast subcluster in the NHC are as follows: *Pdgfd* & *Fgf2* for *Pi16*^+^ fibroblasts; *Cxcl12*, *Mdk*, and *Ccl19* for *Col15a1*^+^ fibroblasts; and *Ccl7* for *Ctla2a*^+^ fibroblasts (**Fig. 6n, Supplementary Fig. 19f–j**). Notably, the *Pi16*^+^ fibroblast subcluster received higher *Igf1* signaling through *Igf1r* than other fibroblasts, notable for its roles in ECM production and invasion.^72, 73^ Together, these findings suggest that *Pi16*^+^ fibroblasts are primarily involved in proliferation, angiogenesis, and tissue remodeling, whereas *Col15a1*^+^ and *Ctla2a*^+^ fibroblasts are preferentially engaged in immunomodulation.

## Discussion

These results directly implicate macrophages as causal in NHC biomaterial-induced tissue remodeling, underscoring their role as a key non-redundant cell type in this process (**Fig. 3a, b**). The NHC’s tissue-remodeling macrophages were linked to pathways involving angiogenesis, ECM remodeling, EMT, promoting inflammation via TNFα/NF-κB, and inducing proliferation via mTORC1 and TrkA (**Fig. 5g, h**). Although both *Ms4a7*^+^ and *Spp1*^+^ macrophages exhibited remodeling-related functions, the *Spp1*^+^ macrophages are especially interesting due to their nearly exclusive presence in the remodeling NHC (**Fig. 5d**). *Spp1*^+^ macrophages were very metabolically active, with nearly all their significantly enriched pathways related to metabolism (**Fig. 5k**). *Spp1*^+^ macrophages are known to influence fibroblast phenotype in fibrosis,^36^ wound healing,^48^ and cancer,^49^ and now for the first time this population has been linked to biomaterial-induced tissue remodeling. Specifically, they communicated through *Spp1*-*Cd44* preferentially with *Pi16*^+^ progenitor-like fibroblasts (**Fig. 5n**). Further, the ability of *Spp1*^+^ macrophages to promote fibroadipo progenitor migration and proliferation^35^ and drive EMT^63^ in other contexts, as well as the known capacity of fibroblasts to convert into adipocytes during wound healing,^74^ represents a potential mechanistic link to long-term outcomes in biomaterial-induced tissue remodeling.

Extensive fibroblast signaling was the most distinctive feature of the remodeling NHC’s inferred communication network (**Fig. 4d**). Their key enriched pathways involved ECM/collagen, angiogenesis, adhesion/motility, and EMT (**Fig. 6h**). The developmental process of EMT, which allows tightly connected epithelial cells to become mobile invasive mesenchymal cells, is especially interesting as these fibroblasts are mesenchymal cells actively infiltrating and remodeling the NHC (**Fig. 3h**). The data suggest fibroblasts played a key role in the observed remodeling, distinct from their known role in the FBR. Despite not being considered immune cells, the NHC’s fibroblasts were extensively involved in immunomodulation, likely functioning as a bridge between the elicited immune response and the resulting tissue remodeling. Specifically, bidirectional macrophage-fibroblast communication involved signaling related to remodeling, immunomodulation, adipogenesis, migration, and angiogenesis. A unique population of *Ctla2a*^+^ fibroblasts was also identified in the remodeling NHC (**Fig. 6g**).

The data demonstrate that NHC biomaterial-induced tissue remodeling progressed through sustained, controlled inflammation. Both predominantly pro-inflammatory and anti-inflammatory markers were significantly increased out to long-term timepoints in the remodeling NHC (**Fig. 2e**). Notably, this phenomenon occurred without any observable adverse physical or behavioral health effects in mice as well as rabbits. Furthermore, the NHC’s significantly enriched pathways included both drivers of inflammation and immune regulatory mechanisms (**Fig. 4b, c**). Though often framed as a negative, inflammation plays a crucial role in this phenomenon. An appropriately balanced host response is crucial, as certain aspects of the immune system support remodeling and others obstruct it. For example, macrophage depletion prevented biomaterial-induced tissue remodeling, but neutrophil depletion accelerated its progress (**Fig. 3a, b**). Taken together, these results demonstrate that controlled and sustained immune activation, with macrophages and fibroblasts as key players, is required for NHC biomaterial-induced tissue remodeling.

The progression of NHC biomaterial-induced tissue remodeling involved the early establishment of a permissive niche which preceded subsequent adipogenesis. This niche was visually apparent as the pre-remodeling zone, which was characterized by its cellular infiltration as well as its unique pattern of collagen deposition, both spatially and via distinct isoforms (**Fig. 1e, g–i**). The thin scaffold-like collagen network throughout the NHC starkly contrasts with the collagen encapsulation observed in typical FBR, as seen with gel controls. This suggests that the body is actively interfacing with the NHC during remodeling, rather than isolating it. Interestingly, preforming this key physical aspect of the niche by replacing the NHC’s poly(ε-caprolactone) nanofibers with collagen nanofibers resulted in substantially faster adipogenesis in a separate study.^1^ Beyond collagen organization, niche establishment was also facilitated by the infiltrating cells’ signaling and effector functions. Notably, macrophage and fibroblast signaling to each other and other cell populations was associated with remodeling as well as immunomodulation (**Fig. 5g–n, Fig. 6h–n**). These cell types were also the primary contributors to adipogenesis-related signaling (**Supplementary Fig. 12b–h**). Intriguingly, at early timepoints they were associated with inhibiting adipogenesis in the ultimately adipogenic NHC material (**Fig. 4g, h**). This suggests early suppression of adipogenesis is part of niche formation, preventing differentiation before the niche is ready to support it. Collectively, these results demonstrate that establishing a pro-remodeling niche is a complex and functionally important stage in NHC’s remodeling.

This study focused on a single biomaterial platform, so generalizability of these findings should be explored in future work. Particularly important areas for further study include additional exploration of key signaling pathways and implicated cell types, as well as identification of the features that bias remodeling toward adipose tissue rather than other soft tissues. More broadly, this work highlights important concepts that may inform the development of future regenerative biomaterials: it suggests that inflammation is nuanced and should not always be viewed as detrimental, that macrophages are critical drivers of tissue remodeling, and that fibroblasts can act as active immunoregulatory partners in shaping host response. Taken together, these findings support the idea that biomaterials may be most effective when they establish an early pro-remodeling niche that enables subsequent tissue remodeling/regeneration.

## Author Contributions

J.L.S., S.K.R., H.-Q.M., and J.C.D. designed experiments, analyzed data, and wrote the manuscript. J.L.S., S.K.R., H.-Q.M., and J.C.D. secured the funding for this study. J.L.S., S.J.B., B.Y.X.N., Z.-C.Y., V.M.Q., J.L.H., K.D.G., R.A.M., and J.C.D. performed experiments. J.L.S., B.L.M., and J.C.D. performed statistical analyses of data sets and aided in the preparation of displays communicating data sets. S.K.R., H.-Q.M., and J.C.D. supervised the study. All authors discussed the results and assisted in the preparation of the manuscript. All authors have given approval to the final version of the manuscript.

## Acknowledgements

This work was supported by LifeSprout Inc. (S.K.R. and H.-Q.M.), the Maryland Stem Cell Research Fund (MSCRFL5375, S.K.R., H.-Q.M., J.C.D.), and the National Institutes of Health through grants R01 DE031488 (S.K.R., H.-Q.M., and J.C.D.), R01 DK135269 (H.-Q.M.), and F31 DE032900 (J.L.S). J.L.S. was also supported by a National Science Foundation Graduate Research Fellowship Program award (DGE-1746891) and an ARCS Foundation Metropolitan Washington Chapter scholar award.

The authors would like to acknowledge and thank the core facilities involved in completing this work. The Johns Hopkins Sidney Kimmel Comprehensive Cancer Oncology Tissue Services, which is supported by the NCI Cancer Center Support Grant (P30 CA006973), processed the histology used in this manuscript. Specifically, Bonnie Gambichler and Miklhail James assisted with these samples. The Johns Hopkins University Research Animal Resources provided animal care and support for this work. Dr. Tyler J. Creamer and Linda Orzolek of the Johns Hopkins Single Cell & Transcriptomics Core provided the scRNA-seq data collection and initial processing, as well as support in planning the sequencing experiment. Dr. Hao Zhang from the Bloomberg Flow Cytometry and Immunology Core was instrumental in assisting with optimizing the cell sorting. Some graphics were created with BioRender.com, as noted in figure legends.

For their personal contributions, we would like to acknowledge Dr. Kavita Krishnan for her guidance on dominoSignal and scRNA-seq, Dr. Kathryn Luly for her instructions on retroorbital injections, Sabrina Chen for her assistance with ChemDraw, as well as Dr. Dan Peng and Dr. Patrick Cahan for their guidance on scRNA-seq annotation and clustering.

## Abbreviations

αSMA: alpha-smooth muscle actin,
Dep: depletion,
DEG: differentially expressed gene,
DGE: differential gene expression,
ECM: extracellular matrix,
EMT: epithelial-to-mesenchymal transition,
FBR: foreign body response,
G’: shear storage modulus,
GSEA: gene set enrichment analysis,
H&E: hematoxylin and eosin,
HA: hyaluronic acid,
HA-Ac: Acrylate-modified hyaluronic acid,
HTO: hashtag oligonucleotide,
KO: knockout,
Macro: macrophage,
MAL: maleimide,
NHC: nanofiber-hydrogel composite,
Neut: neutrophil,
NK: natural killer,
PBS: phosphate buffered saline,
PCL: poly(ε-caprolactone),
PCL-MAL: maleimide-functionalized poly(ε-caprolactone),
PEG-SH: dithiol poly(ethylene glycol),
qPCR: quantitative polymerase chain reaction,
RT: reverse transcription,
scRNA-seq: single-cell RNA sequencing,
SEM: standard error of the mean,
TCR: T-cell receptor,
UMAP: Uniform Manifold Approximation and Projection,
WT: wild type

## Methods

### Materials Synthesis

Acrylate-modified hyaluronic acid (HA-Ac) was made by mixing 1% sodium hyaluronate (HA; 1.5 × 10^6^ Da; LifeCore Biomedical, Chaska, MN) in phosphate buffered saline (PBS) (pH 8.5) with glycidyl acrylate (TCI, Portland, OR) at a 100:3 ratio (v/v) for 16 h at 37 °C on a magnetic stirrer. After precipitation of the HA-Ac by adding the reaction mixture to ethanol at a 1:10 volume ratio, the precipitate was washed three times with ethanol and acetone and dehydrated with compressed air. It was then redissolved in PBS (pH 7.4) and stored at 4 °C until use.

The poly(ε-caprolactone) (PCL) nanofiber fragments were made according to previously published protocols.^25, 75^ In brief, PCL solution (16%, w/w) was made by dissolving PCL in a 9:1 (v/v) dichloromethane and dimethylformamide mixture. This solution was electrospun into PCL nanofibers which were then surface activated with plasma treatment to generate carboxylic functional groups. The carboxyl groups were activated and converted to maleimide (MAL) groups, generating MAL-functionalized PCL (PCL-MAL) fibers. This was done by treating the fibers with ethyl dimethylaminopropyl carbodiimide and N-hydroxysuccinimide at a molar ratio of 1:4:4, respectively. N-(2-Aminoethyl) MAL was added at a molar ratio of carboxyl groups to amine groups of 1:2, and the mixture was gently shaken to facilitate this conversion. A cryogenic mill (Freezer/Mill 6770, SPEX SamplePrep, Metuchen, NJ) was used to break the PCL-MAL fibers into fragments, which averaged in length from 20-100 μm. PCL-MAL fibers were sterilized in three cycles of 70% (v/v) ethanol followed by distilled water, then lyophilized and stored at -20 °C until use.

NHCs were generated by mixing HA-Ac, PCL-MAL, and dithiol poly(ethylene glycol) (PEG-SH) (molecular weight: 5 kDa, JenKem Technology, Plano, TX) at 37 °C for 16 h to crosslink them. The composites were formulated with PCL-MAL fibers (30 mg/mL) and varied concentrations of HA-Ac (7 or 12 mg/mL) in PBS (pH 7.4). The concentration of PEG-SH was selected to keep the thiol-acrylate ratio at 1.0. The PEG-SH concentrations used were 4.80 and 7.65 mg/mL, respectively. HA control materials were generated using the same method, except without the addition of PCL-MAL fibers. HA-Ac concentrations used were 10 and 17 mg/mL, and the corresponding PEG-SH concentrations used were 5.75 and 9.80 mg/mL. After gelation, materials were mechanically fragmented to enhance their injectability. Materials were passed 6 times through a stainless-steel mesh screen with an opening size of 0.003 inches, and stored at 4 °C until use.

### Materials Synthesis (Commercial Grade)

For the commercial-grade NHC (Lumina), a slightly modified version of the previously described NHC synthesis was carried out. Briefly, 50 kDa PCL (Corbion, Netherlands) was formed into fiber sheets by electrospinning (Ucalery, China) at a feed rate of 0.273 mm/min at a concentration of 12.6% w/w of polymer in 2,2,2-trifluoroethanol (Sigma, USA). The positive voltage was 15.6 kV, with a negative voltage of 3.5 kV applied to a rotating drum target. The fiber sheets were vacuum dried, then plasma treated for 20 min per side to activate the fiber surface (Harrick PDC-001, Harrick, USA). The activated fiber surface was modified with maleimide groups by immersion in a buffer of 0.1 M MES (2-(N-morpholino)ethanesulfonic acid, Sigma) (pH 6) mixed with ethanol (50:50, v/v) containing 10 mg/mL EDC, 10 mg/mL NHS, and 10 mg/mL N-(2-aminoethyl)maleimide (final concentration) for 2 h. The PCL fiber mats are disrupted into individual fiber fragments by cryogenic grinding (Freezer/Mill 6770, Spex SamplePrep, USA) and high shear mixing (Silverson L5M-A High Shear Mixer, USA) before being vacuum dried.

Hyaluronic acid was modified with glycidyl acrylate (BOC Sciences, China) so that approximately 10% of the disaccharide subunits contain an acrylate group. Polyethylene glycol dithiol (5,000 Daltons, Jenkem USA) was used to crosslink the gel composite by reacting with the acrylate groups on the hyaluronic acid and the maleimide groups on the PCL fiber surfaces through a Michael addition reaction without any additives or byproducts. The gel components were separately sterilized (HA-Acrylate and PEG-SH stock solutions through sterile filtration and the PCL fibers are sterilized through E-beam irradiation), then aseptically mixed and incubated overnight at 37 °C to gel. The gel was formulated to be comprised of 10 mg/mL HA-Ac, 15 mg/mL PEG-SH, 30 mg/mL of PCL fibers, and 3 mg/mL lidocaine hydrochloride in phosphate-buffered saline. The bulk gel was then broken up into gel fragments that are approximately 150 µm in diameter by being pressed through a series of screens (with decreasing pore sizes of 500, 250, and 150 µm) before being loaded into syringes.

### Mouse Animal Experiments

All murine animal work was carried out under a research protocol approved by the Johns Hopkins University Animal Care and Use Committee. All mice were male and 6-8 weeks of age. Unless otherwise noted, C57BL/6J mice (strain 000664, The Jackson Laboratory) were used. NHC or gel control materials were injected subcutaneously onto the backs of mice, with each mouse receiving no more than 6 injections. All injections were 100 μL in volume and done with a 25-gauge needle (Cat# 305122, BD) while animals were under isoflurane. Mice were not shaved to prevent them from scratching the area. Animals were carefully monitored after injections until they were awake and moving around normally.

### Mouse Depletion and Knockout Models

The following strains of mice were used for knockout experiments: muMt- mice (strain 002288, The Jackson Laboratory), B6 nude mice (strain 000819, The Jackson Laboratory), Rag2/Il2rg double knockout mice (model 4111, Taconic Biosciences), Il15 KO mice (strain 034239, The Jackson Laboratory). For depletion experiments, C57BL/6J mice (strain 000664, The Jackson Laboratory) were injected with depletion agents. To ensure depletion before materials were injected, all depletions were started at day -3 and material injections occurred at day 0. For macrophage depletion, 200 μL of Clodrosome (CLD-8909, Encapsula NanoSciences) per mouse was injected every 3 days. Of the 200 μL, 100 μL was delivered via retroorbital injection and 100 μL was delivered via subcutaneous injection at the site of the materials. As a note, Clodrosome administration leaves mice highly susceptible to opportunistic infections so extra care must be taken when handling these animals and all of them that begin the treatment will not make it to the end. For natural killer cell depletion, 50 μL of an anti-asialo GM1 antibody (986-10001, FUJIFILM Wako Pure Chemical Corporation) diluted with 100 μL of sterile cell culture grade water per mouse was injected intraperitoneally every 6 days. For neutrophil depletion, an anti-Ly6g (clone 1A8) antibody (Cat # 127650, BioLegend) was diluted with 1x PBS to inject 250 μg of antibody in 150 μL total volume per mouse injected intraperitoneally every 3 days.

### Rabbit Experiments

Rabbit implantation, tissue collection, and histology were conducted by North American Science Associates (NAMSA). The rabbit implantation study was modeled upon the ISO 10993-6 biocompatibility testing standard. Young adult female New Zealand White rabbits (Oryctolagus cuniculus) 2.5-3.5 kg at selection were purchased from Robinson Services Inc. and housed per NAMSA standard operating procedures. For each implantation site, a location marker (10 mm × 1 mm × 1 mm plug of sterilized high-density polyethylene) was implanted via a 16 G needle into the subcutaneous tissue of the rabbit dorsal region, then 0.4 cc of the test article was injected. All groups were implanted into each animal. The test and control sites were excised as full thickness skin sections that included a tissue margin surrounding each site. Tissues were fixed in 10% neutral buffered formalin, trimmed, and embedded in paraffin per standard histological techniques. The blocks were sectioned at approximately 4 to 5 µm and stained with H&E.

Lumina was the test article of interest, Radiesse (Merz) was used as a commercial control article, and the negative control was 0.9% NaCl. Radiesse Lidocaine injectable implant is an opaque, sterile, nonpyrogenic, semi-solid, cohesive implant, whose principal component is synthetic calcium hydroxyapatite suspended in a gel carrier of glycerin, sodium carboxymethylcellulose, 0.3% lidocaine hydrochloride, and sterile water for injection. Radiesse is FDA-approved for sub-dermal implantation in the face and hands.

### Rheology

The shear storage modulus (G’) of the materials was measured with a rheometer (ARG2, TA Instruments) using a smooth parallel plate geometry of 8 mm. Samples were injected into the sample holder, then shaped into an 8 mm disk. The linear viscoelastic region was determined using a strain sweep (0.01 – 10% strain), then a strain within the linear range was chosen to run a frequency sweep (0.1 – 10 Hz). Measurements were taken from distinct samples.

### Scanning Electron Microscopy

The materials’ microstructures were analyzed with the following procedure. Firstly, gel control and NHC materials were lyophilized, followed by imaging with a scanning electron microscope (Thermo Scientific Helios G4 UC, USA). Precursor solutions were cast into an 8 mm polydimethylsiloxane mold and set to crosslink for 16 h. Then, the gel control or NHC materials were lyophilized at -20 °C for 24 h. Cross sections of the gel control and NHC materials were imaged using a scanning electron microscope at an accelerating voltage of 5 kV.

### Histology

Materials were explanted with skin attached and placed into 4% PFA for at least 48 h for fixation. After fixation, the samples were submitted to the Johns Hopkins Oncology Tissue and Imaging Services Core facility for further processing, embedding, sectioning, and staining. Briefly, samples were dehydrated, embedded in paraffin wax, and sectioned into 5 µm slices. Cross sections down the middle of materials were obtained. H&E staining as well as Masson’s Trichrome staining was performed. Percent area adipose, pre-remodeling zone, and collagen were assessed using the mouse Masson’s Trichrome images with Fiji/ImageJ. Cellular infiltration was assessed using the mouse H&E images with Fiji/ImageJ. For rabbit data, H&E images were used for all quantification with Fiji/ImageJ.

### Immunofluorescence

Upon collection, samples were frozen fresh in Scigen O.C.T. compound cryostat embedding medium (Cat# 23-730-625, Fisher Scientific). OCT embedded samples were sectioned to be 8 µm thick (Leica CM3050 S Cryostat). Samples were thawed (5 min), circled with a PAP pen (Cat# Z672548, Millipore Sigma) and rehydrated in PBS (5 min). Then, they were fixed with 4% PFA (20 min), washed in PBS (3 × 5 min), and permeabilized on ice with 1% Triton X-100 (Cat# T8787, Millipore Sigma) (5 min). Blocking was done with Dako buffer (Cat # S302283-2, Agilent) (1 h), then staining with primary antibodies (**Table 1**) was carried out (2 h, room temperature). After washing slides with PBS (3 × 5 min), they were mounted with coverslips using ProLong Diamond Antifade Mountant (Cat# P36961, ThermoFisher) and were left to cure for 24 h before being sealed with clear nail polish (Cat# 72180, Electron Microscopy Services).

**Table 1.** Antibodies Used for Immunofluorescence Microscopy.

| Target | Clone | Fluor | Manufacturer | Cat # | Dilution |
| --- | --- | --- | --- | --- | --- |
| NucBlue | - | - | Invitrogen, ThermoFisher | R37606 | 2 drops per 1 mL |
| αSMA | 1A4 | Cy3 | Millipore Sigma | C6198 (100 μL) | 1:200 |
| CD68 | FA-11 | AF647 | BioLegend | 137004 (100 μg) | 1:200 |

### Flow Cytometry

Materials were explanted, minced, and enzymatically digested with 7 mg/mL hyaluronidase type I-S (Cat# H3506, Millipore Sigma), 7 mg/mL hyaluronidase type II (Cat# H2126, Millipore Sigma), and 2 mg/mL collagenase D (Cat# 11088858001, Millipore Sigma) in serum free media (Cat# 61870-036, Gibco) for 1 h on a shaker in an incubator at 37° C. Samples were processed on a gentleMACS Dissociator (Miltenyi Biotec) in prepared PEB buffer (PBS, 0.5% BSA, 2 mM EDTA). They were subsequently filtered with a 40 µm mesh, spun down, and had RBC lysis performed if necessary (Cat# 00-4333, 1× RBC Lysis Buffer, eBioscience. Samples were Fc blocked (Cat# 14-0161-81, ThermoFisher) then stained with properly titrated antibodies (**Table 2**) in Flow Cytometry Staining Buffer (Cat# 00-4222, eBioscience). Super Bright Complete Staining Buffer (SB-4401-75, ThermoFisher) was added, as more than one polymer dye-conjugated antibody was used in the antibody panels. Cells were then washed. If intracellular staining was required, cells were fixed for 30 min with 1% PFA (Cat# 15714-S, Electron Microscopy Sciences) in PBS and permeabilized for 5 min with 0.1% Triton X-100 (Cat# T8787, Millipore Sigma). If intranuclear staining was required, the Foxp3/Transcription Factor Staining Buffer Set (Cat# 00-5523-00, eBioscience/ThermoFisher) was used instead. After washing, intracellular or intranuclear staining was performed. After washing, samples were run on a flow cytometer (Attune NXT, ThermoFisher). FCS files were analyzed using FlowJo software.

**Table 2.**
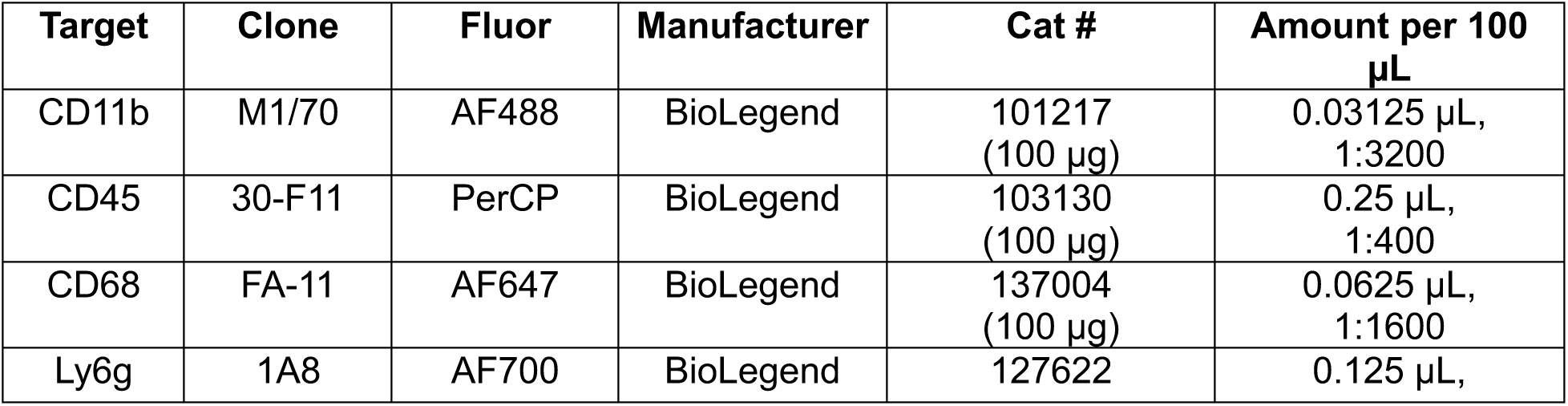
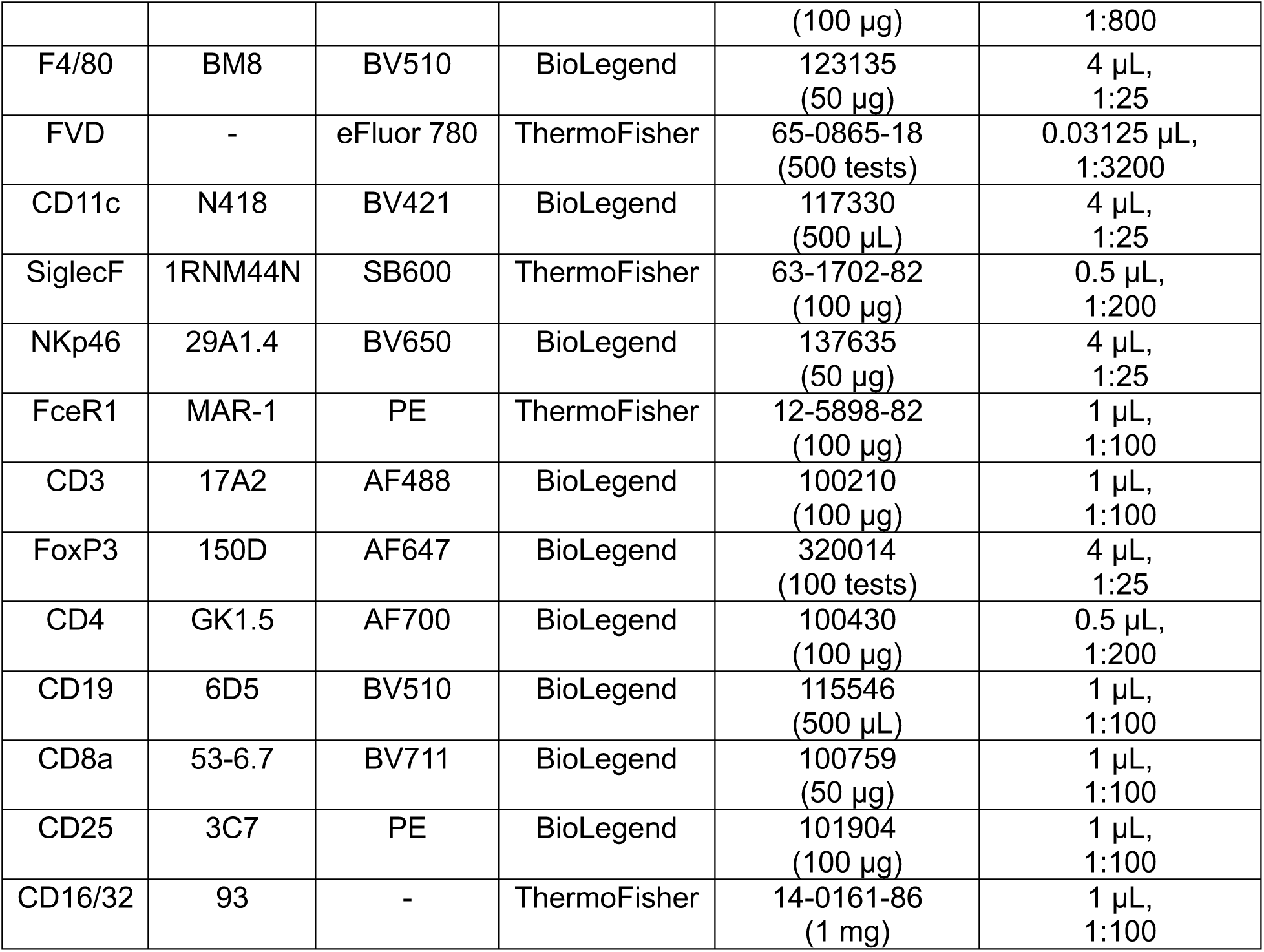
Antibodies Used for Flow Cytometry.

### RNA Isolation & RT-qPCR

RNA was isolated with TRIzol (Cat#15596018, ThermoFisher) & a power homogenizer (Polytron/VWR) from samples snap-frozen in liquid nitrogen. In brief, after samples were homogenized thoroughly, RNA was precipitated with chloroform, washed twice with 75% ethanol, dried, and then resuspended in DEPC-treated water (Cat# 46-2224, Invitrogen). Glycogen (Cat#R0561, ThermoFisher) was added during the precipitation step for small samples. The RNA concentration and quality were verified using a NanoDrop 2000 spectrophotometer (ThermoFisher).

For DNAse and reverse transcription (RT), 1 μg total RNA was used for each sample in a uniform volume of DEPC-treated water (Cat# 46-2224, Invitrogen). For DNAse, 10× RT buffer and dNTP mix from the High-Capacity cDNA Reverse Transcription Kit (Cat #4368814, Applied Biosystems) as well as DNAse (Cat# M6101, Promega), and RNAse inhibitor (Cat# N8080119, ThermoFisher) were added to each sample. A DNAse cycle (1 h at 37 °C, 10 min at 75 °C, and an infinite hold at 4 °C) was performed on a ThermoCycler to remove potential contaminating DNA.

Afterwards, for RT, samples had 10× RT random primers and reverse transcriptase enzyme from the High-Capacity cDNA Reverse Transcription Kit (Cat #4368814, Applied Biosystems) added. They were then placed back on the ThermoCycler for an RT cycle (10 min at 25 °C, 2 h at 37 °C, 5 min at 85 °C, infinite hold at 4 °C) to reverse transcribe the RNA into cDNA. The cDNA samples were diluted 1:20 with 50 μg/mL yeast RNA (Cat# AM7118, Invitrogen), which was previously diluted in Milli-Q water. At this point, samples were pooled if desired.

The cDNA samples were combined with Power SYBR Green (Cat# 4367659, Applied Biosystems), forward and reverse primers (**Table 3**) (Eurofins Genomics), and Milli-Q water, then run in triplicate on a QuantStudio 5 Real-Time PCR System (ThermoFisher). Samples were incubated for 2 min at 50 °C, then 10 min at 95 °C, followed by 40 cycles of 95 °C for 15 sec and 60 °C for 60 sec. Results were analyzed using the comparative CT (ΔΔCT) method, which employs a housekeeping gene, and are presented as RNA levels relative to controls, as indicated.

**Table 3.**
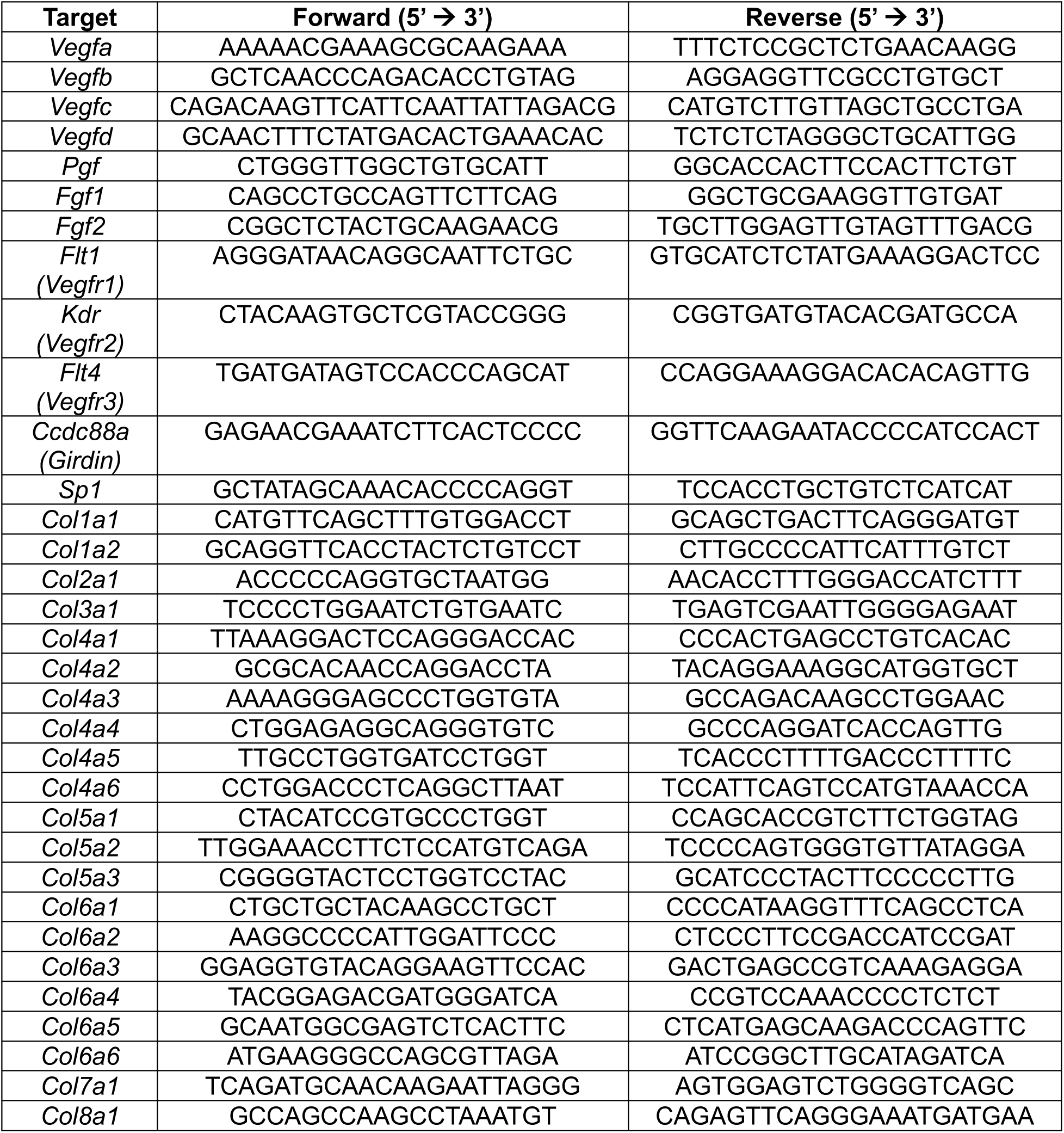
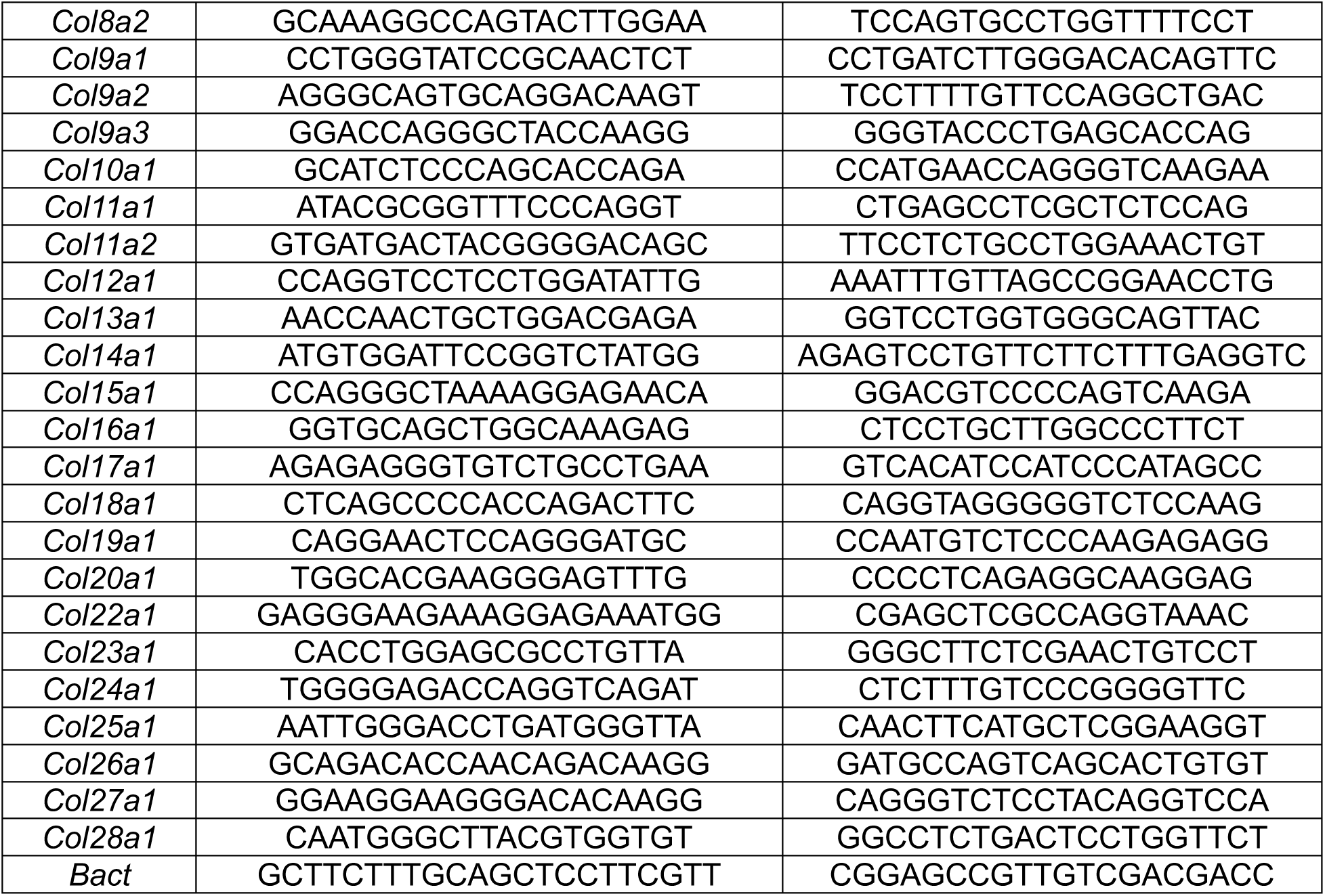
Sequences of Forward and Reverse Primers Used for qPCR.

### NanoString

RNA was isolated from snap-frozen samples and quantified as described above. Samples were brought to an appropriate concentration (100 ng/μL), and 500 ng of each sample was processed according to the manufacturer’s protocols from NanoString. Briefly, samples were hybridized for 16-48 h, cartridges were prepared using the nCounter Prep Station, and subsequently run on the nCounter Digital Analyzer. Samples were run with a custom 120-gene panel for mouse to obtain absolute copy numbers of RNA. Data were processed using the nSolver 4.0 and nSolver Advanced Analysis 2.0.134 software. nSolver was used for data normalization and nSolver Advanced Analysis differential expression analysis was used to calculate *p*-values. The Benjamini-Hochberg method of estimating false discovery rates was used to perform *p*-value adjustment.

### Statistics

Statistical significance was calculated using GraphPad Prism (Version 10) for all assays except NanoString and scRNA-seq, which were analyzed with nSolver Advanced Analysis and in R, respectively, with details provided in their respective methods sections. Specific statistical tests and replicate numbers are detailed in the figure legends. Data is shown as mean ± SEM with 3 to 5 replicates per treatment group, unless otherwise indicated. Comparisons between groups were performed using a two-way ANOVA followed by Tukey’s multiple comparisons test, unless otherwise indicated. Comparisons were considered significant when *p* < 0.05 (\**p* < 0.05, \*\**p* < 0.01, \*\*\**p* < 0.001, \*\*\*\**p* < 0.0001).

### Single-Cell RNA Sequencing

Materials were explanted, minced, and enzymatically digested as described in the flow cytometry section. RBC lysis was not performed. Samples were then Fc blocked with TruStain FcX PLUS (Cat# 156603, BioLegend) in cell staining buffer (Cat# 420201, BioLegend) for 10 min at 4 °C. Then, samples were barcoded by adding titrated BioLegend TotalSeq-C Hashtag reagents (Cat# 155861, 155863, 155865, 155867, & 155869, BioLegend) at 0.125 μg per sample and incubating for 30 min at 4 °C. Samples were washed 3 times, filtered, and stained with Calcein AM (Cat# C3100MP, Invitrogen), Vybrant DyeCycle Violet Ready Flow Reagent (Cat# R37172, Invitrogen), and propidium iodide (Cat# P3566, Invitrogen). Live cells were then sorted by the Johns Hopkins Bloomberg Flow Cytometry, Immunology, and Cell Sorting Core (Beckman Coulter - MoFlo XDP) to separate them from the fibers that could clog the sequencing machine. Live cells were sorted into media (Cat# 61870-036, Gibco) with 2% BSA (Cat# A3294, Sigma) added and then taken to the Johns Hopkins Single Cell & Transcriptomics Core for 5’ single-cell RNA sequencing.

Cell counts and viabilities were determined using the Cell Countess 3 with Trypan blue staining. Cells were combined, with three or four biological replicates per group (n = 3 for day 7 NHC, day 7 gel, day 35 NHC; n = 4 for day 35 gel), at desired ratios to capture target cell number based on a 65% capture rate. Cells were combined with RT reagents and loaded onto 10X Next GEM Chip N along with 5’ HT gel beads. The NextGEM protocol was run on the 10X Chromium X to create GEMs (gel bead in emulsion), composed of a single cell, gel bead with unique barcode and UMI primer, and RT reagents. Approximately 180 μL of emulsion was retrieved from the chip, split into two wells, and incubated (45 min at 53 °C, 5 min at 85 °C, then cooled to 4 °C), generating barcoded cDNA from each cell. The GEMs were broken using Recovery Agent, and cDNA was cleaned, following the manufacturer’s instructions using MyOne SILANE beads. cDNA was amplified using Feature cDNA primers 4 (45 sec at 98 °C; 11 cycles: 20 sec at 98 °C, 30 sec at 63 °C, 1 min at 72 °C; 1 min at 72 °C; then cooled to 4 °C). Samples were cleaned using 0.6X SPRIselect beads. Supernatant containing the hashing barcode tag was reserved and cleaned with 2.0X SPRI reagent. Quality control assays were completed using Qubit and Bioanalyzer to determine size and concentrations. Twenty μL of amplified cDNA were used in library preparation. Fragmentation, end repair, and A-tailing were completed (5 min at 32 °C, 30 min at 65 °C, then cooled to 4 °C), and samples were cleaned up using double-sided size selection (0.6X, 0.8X) with SPRIselect beads. Adaptor ligation (15 min at 20 °C, then cooled to 4 °C), 0.8X cleanup, and amplification were performed, with PCR using unique i7 and i5 index sequences. Libraries underwent a final cleanup using double sided size selection (0.6X, 0.8X) with SPRIselect beads. HTO tag cDNA was amplified with PCR using unique i7 and i5 index sequences. Library quality control was performed using Qubit, Bioanalyzer, and KAPA library quantification qPCR kit. Libraries were sequenced on the Illumina NovaSeq 6000 using v1.5 kits, targeting 50,000 reads/cell for gene expression and 5,000 reads/cell for VDJ and CITE-seq, at read lengths of 28 (R1), 10 (i7), 10 (i5), 91 (R2).

### Single-Cell RNA Sequencing Computational Processing

Demultiplexing and FASTQ generation was completed using Illumina’s BaseSpace software. FASTQ files were processed using Cell Ranger Multi v7.1.0 (10X Genomics), which aligned reads to the reference genome (mm10-2020-A), assigned unique molecular identifiers to generate gene expression matrices, and performed hashtag oligonucleotide demultiplexing to assign cells to their respective samples.

Demultiplexed data were processed using R via RStudio. Seurat v5.0.3 was used for filtering, normalization, dimensional reduction, integration, clustering, and annotation.^76^ Data was filtered based on unique molecular identifier counts (>500, <50,000), number of features (>350, <6,000), and percentage of mitochondrial reads (>5%). Normalization and variance stabilization were performed using sctransform.^77, 78^ Dimensional reduction was done using principal component analysis, and 50 dimensions were selected for downstream analysis based on visualization with an elbow plot. Data was integrated using Harmony to correct for batch effects.^79^ A Shared Nearest Neighbor graph was constructed, and Louvain clustering was performed. Clustering resolution was determined based on biologically relevant markers and silhouette scores. Uniform Manifold Approximation and Projection (UMAP) was used for data visualization, and clusters were annotated based on DEGs and canonical markers. Clusters of interest were isolated and further analyzed to identify subclusters, following the same pipeline described above.

Differential abundance analysis was performed using edgeR v4.0.16’s quasi-likelihood generalized linear model framework with non-trended dispersion estimates to test for significant changes in per-cluster cell abundance.^80^ For pseudobulk DGE analysis, samples were aggregated, DEGs were identified using DESeq2^81^ v1.42.1, and log2 fold change shrinkage was performed via the apeglm method.^82^ Pseudobulk DGE comparisons were performed between NHC and gel control groups, overall and by cluster. For clusters with insufficient sample size for pseudobulk analysis, DEGs were calculated using the Seurat implementation of the Wilcoxon rank-sum test. Wilcoxon comparisons were performed by comparing the cluster of interest to all other cells in the entire dataset. The fgsea package^83^ v1.28.0 was utilized to perform rank-based GSEA using the Hallmark, BioCarta, Reactome, and WikiPathways collections of gene sets, with DGE results ranked by -log(*p*-value)*sign(log2FoldChange). Adjusted *p*-values were calculated using the Benjamini-Hochberg method to control the false discovery rate. Additional adipogenesis pathways were sourced from MSigDB’s curated gene sets. All shown Module Scores were calculated using the Seurat “AddModuleScore” function on the relevant subset of cells shown in each figure. Genes most correlated to pathways were selected by identifying the top 150 genes most correlated in expression to the corresponding MSigDB pathway (by Pearson Correlation between the gene’s SCT assay value and the pathway’s Seurat-calculated ModuleScore), excluding genes in the corresponding MSigDB pathway. To compare module scores, per-cell module scores were averaged within each sample to generate pseudobulk estimates. Pairwise differences between groups were assessed using t-tests between sample-level means. Adjusted *p*-values were calculated using the Benjamini-Hochberg method to control the false discovery rate. Omnibus comparisons were assessed using ANOVA followed by Tukey’s multiple comparisons test.

Cell signaling was analyzed using dominoSignal v1.0.3, with further analysis leveraging the “network_to_df” function in the developmental branch “Krishnan_network2df”, commit db1df06.^84,85^ This dominoSignal analysis was run on clusters with >150 cells per group, excluding cycling cells and doublets, and it was run a second time with the fibroblasts, macrophages, and neutrophils split into their respective subclusters. Transcription factor activation scores were calculated using pySCENIC^86^ (Aerts Lab Docker image aertslab/pyscenic:0.12.1), and cellphoneDB^87^ v4.0.0 was used to map ligand-receptor linkages. Ligand signaling score is a metric created to quantify outgoing signaling per ligand. It is calculated by multiplying a ligand’s gene expression level by the corresponding number of dominoSignal ligand-receptor linkages between the sending and receiving cluster including that ligand, and log normalizing. Pseudotime trajectory analysis was performed using Monocle3 (v1.4.26). Root cells were defined as macrophage and monocyte cells ranking in the top 1% by Seurat-calculated module score for monocyte-derived lineage markers (genes *Myb*, *Ms4a3*, *Ccr2*, *Ly6c2*, & *Sell*), restricted to those mapping to the largest connected component among principal-graph vertices associated with these high-scoring cells.

## Data Availability

All data supporting the findings of this study are available within the article and its supplementary files. Reasonable requests for additional information will be fulfilled by the corresponding authors. The sequence raw data are available in the NCBI Gene Expression Omnibus (GEO) under accession number GSE300761.

