## Supplemental Materials for "Fibroblast signaling influences macrophage-dependent, biomaterial-induced tissue remodeling"

**Supplemental Index of Sections**

1) Supplementary Figures 3

Supplementary Figure 1. NHC material characterization and stiffness comparisons.

Supplementary Figure 2. Extended representative histology.

Supplementary Figure 3. Quantification of Col6a3 expression with NanoString and qPCR.

Supplementary Figure 4. NHC-induced remodeling in New Zealand White rabbits.

Supplementary Figure 5. Extended representative histology.

Supplementary Figure 6. Flow cytometry gating for lymphocyte panel.

Supplementary Figure 7. Flow cytometry gating for myeloid panel.

Supplementary Figure 8. Full NanoString heatmap of all genes for WT mice.

Supplementary Figure 9. Additional knockout/depletion data.

Supplementary Figure 10. Additional scRNA-seq identity markers and neutrophil information.

Supplementary Figure 11. Overall communication networks showing selected subclusters of interest.

Supplementary Figure 12. Additional adipogenesis pathways.

Supplementary Figure 13. NHC materials influence early adipogenesis-related signaling during niche establishment.

Supplementary Figure 14. Clustering and subclustering of scRNA-seq data.

Supplementary Figure 15. Exclusion of low-quality macrophages.

Supplementary Figure 16. Macrophage trajectory inference reveals skewed populations by material type.

Supplementary Figure 17. Extended scRNA-seq analysis for macrophages.

Supplementary Figure 18. Extended scRNA-seq differential gene expression and gene set enrichment analysis for fibroblasts.

Supplementary Figure 19. Extended scRNA-seq cell signaling analysis for fibroblasts.

2) Supplementary Table

Supplementary Table 1. Flow Cytometry Cell Population Abundance. 27

### Supplementary Figures

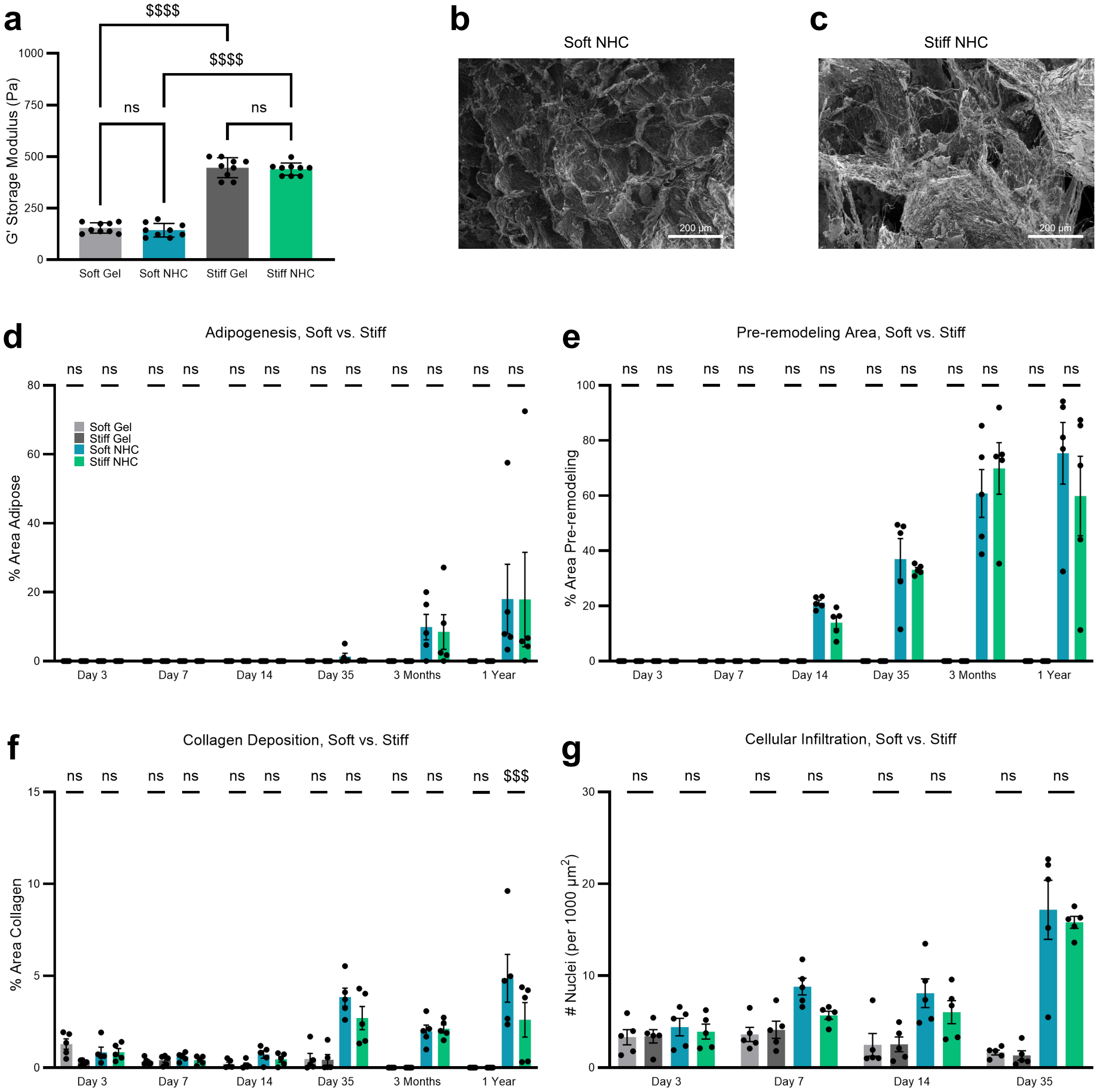


**Supplementary Figure 1. NHC material characterization and stiffness comparisons. a,** Rheology demonstrating stiffness-matching of NHC and gel control materials (soft: 150 Pa or stiff: 450 Pa). Data, mean ± SEM (n = 9 replicates). One-way ANOVA followed by Tukey's multiple comparisons test. **b, c,** SEM image showing structure of soft NHC (**b**) and stiff NHC (**c**). Scale bars are as annotated. **d–f,** Analysis of Masson's Trichrome histological images quantifying the amount of adipogenesis (**d**), pre-remodeling zone (**e**), and collagen deposition (**f**) inside the materials at various timepoints. Plots show the percent area. Data, mean ± SEM (n = 5 biological replicates). Two-way ANOVA followed by Tukey's multiple comparisons test. **g,** Analysis of H&E histological images quantifying the amount of cellular infiltration inside the materials at various timepoints. Plot shows the nuclei per area. Data, mean ± SEM (n = 5 biological replicates). Two-way ANOVA followed by Tukey's multiple comparisons test. In this figure's graphs, statistics shown are ($^{\$}$) timepoint and material-matched comparisons of soft vs. stiff. $^{\$}p < 0.05$; $^{\$\$}p < 0.01$; $^{\$\$\$}p < 0.001$; $^{\$\$\$\$}p < 0.0001$.

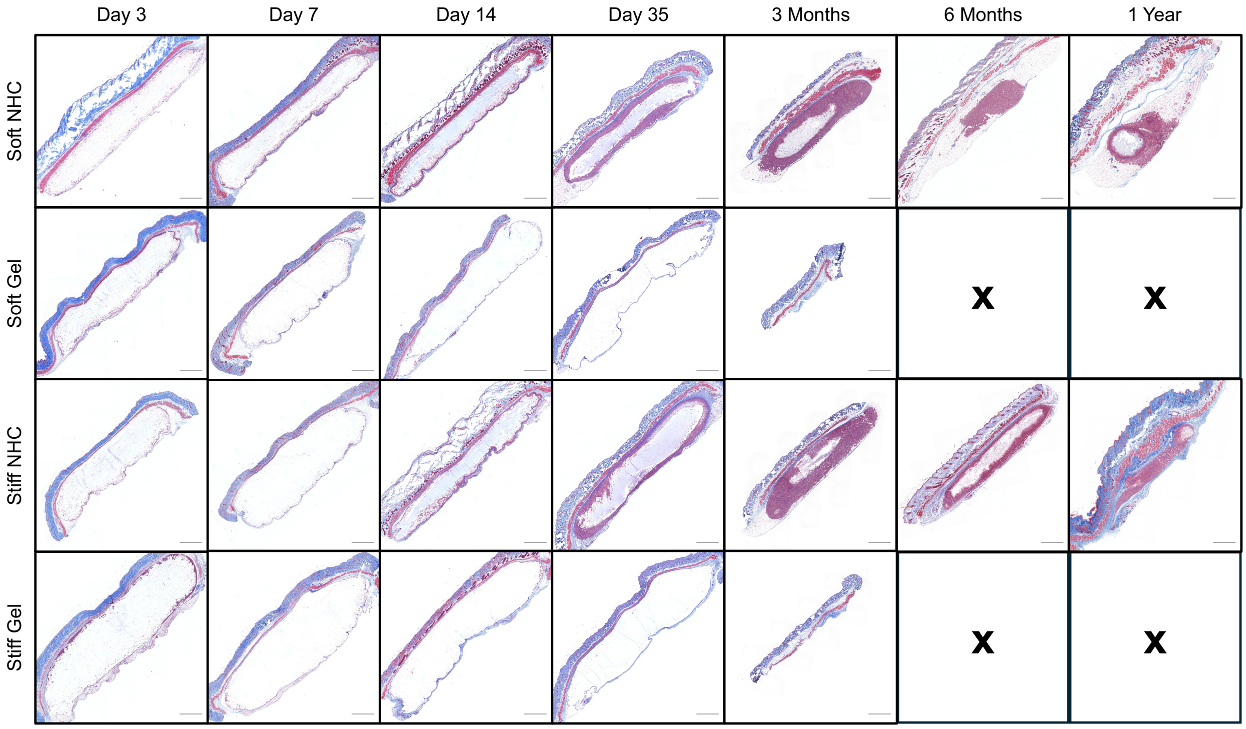


**Supplementary Figure 2. Extended representative histology.** Representative Masson's Trichrome-stained histologic cross-sections of materials subcutaneously injected into C57BL/6 mice, showing gross morphology including skin. Soft materials for day 7, day 35, 3 months, and 1 year also appear in Figure 1d. Scale bars are 1000 µm.

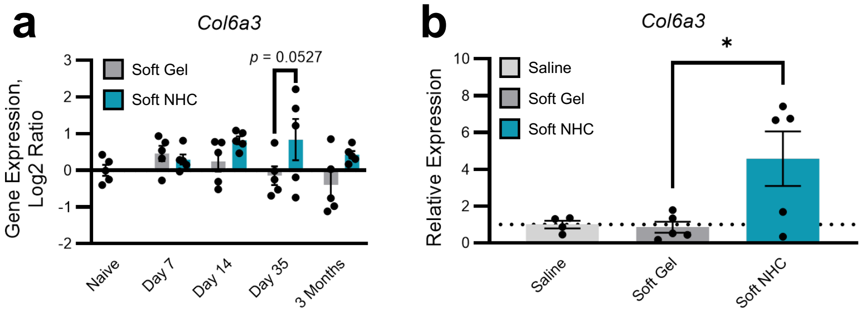


**Supplementary Figure 3. Quantification of *Col6a3* expression with NanoString and qPCR.** **a,** NanoString gene expression for *Col6a3* on materials explanted from C57BL/6 mice at various timepoints. Plot shows the absolute transcript counts normalized to naïve fascia (scale: log2 ratio). Data, mean ± SEM (n = 5 biological replicates). Two-way ANOVA followed by Šídák's multiple comparisons test. **b,** qPCR gene expression analysis for *Col6a3* on materials explanted from C57BL/6 mice at day 35. Plots show the relative expression normalized to saline control. Data, mean ± SEM (n = 5 biological replicates). One-way ANOVA followed by Šídák's multiple comparisons test. This was done to confirm significance for *Col6a3*, as the NanoString *p*-value was just over the cutoff. In this figure's graphs, statistics shown are (*) timepoint and stiffness-matched comparisons of NHC vs. gel control. *$p < 0.05$, **$p < 0.01$, ***$p < 0.001$, ****$p < 0.0001$.

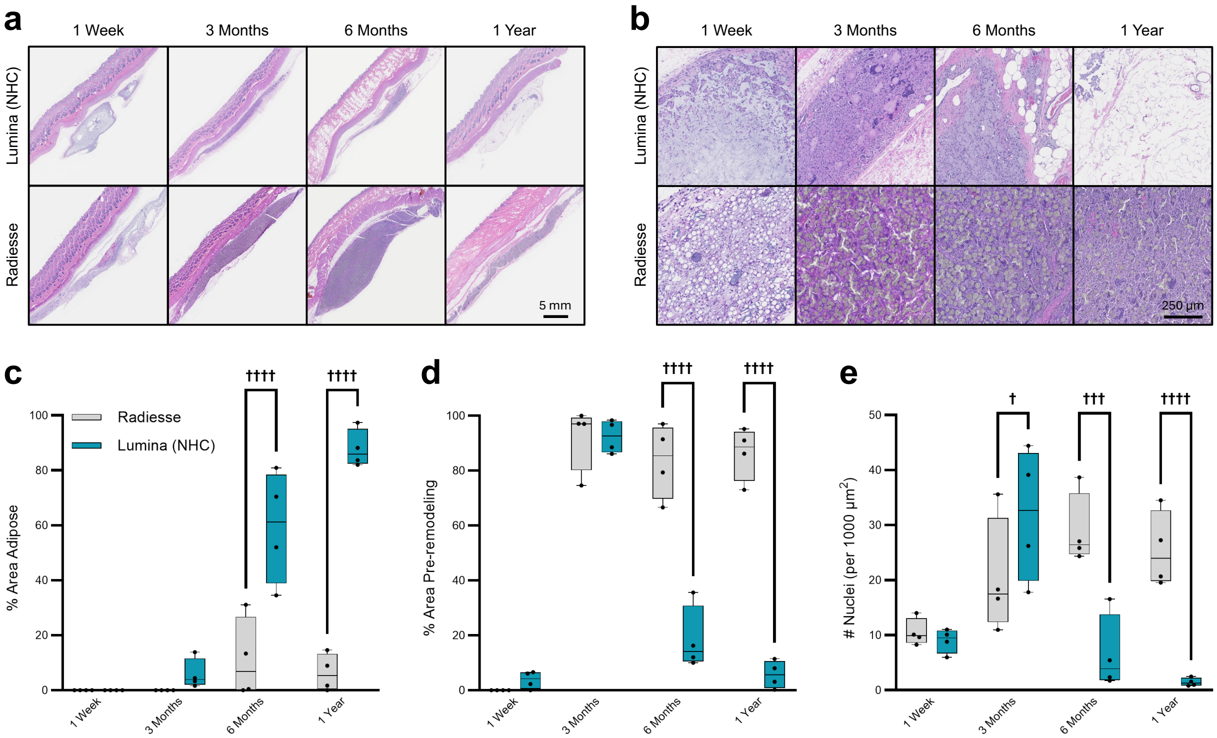


**Supplementary Figure 4. NHC-induced remodeling in New Zealand White rabbits. a, b,** Representative H&E-stained histologic cross-sections of commercial-grade NHC materials (Lumina) and a control commercially available dermal filler (Radiesse) that were subcutaneously injected into New Zealand White rabbits. Overall zoomed-out images showing gross morphology including skin with 5 mm scale bar (**a**) and zoomed-in view inside materials with 250 µm scale bar (**b**) are shown. **c–e,** Analysis of H&E histological images quantifying the amount of adipogenesis (**c**), pre-remodeling zone (**d**), and cellular infiltration (**e**) inside the materials at various timepoints. Plots show the percent area of the component of interest from the total area of the biomaterial. Data, min to max box and whisker (n = 4 biological replicates). Two-way ANOVA followed by Tukey's multiple comparisons test. In this figure's graphs, statistics shown are (†) timepoint-matched comparisons of Lumina vs. Radiesse. $^{\dagger}p < 0.05$, $^{\dagger\dagger}p < 0.01$, $^{\dagger\dagger\dagger}p < 0.001$, $^{\dagger\dagger\dagger\dagger}p < 0.0001$.

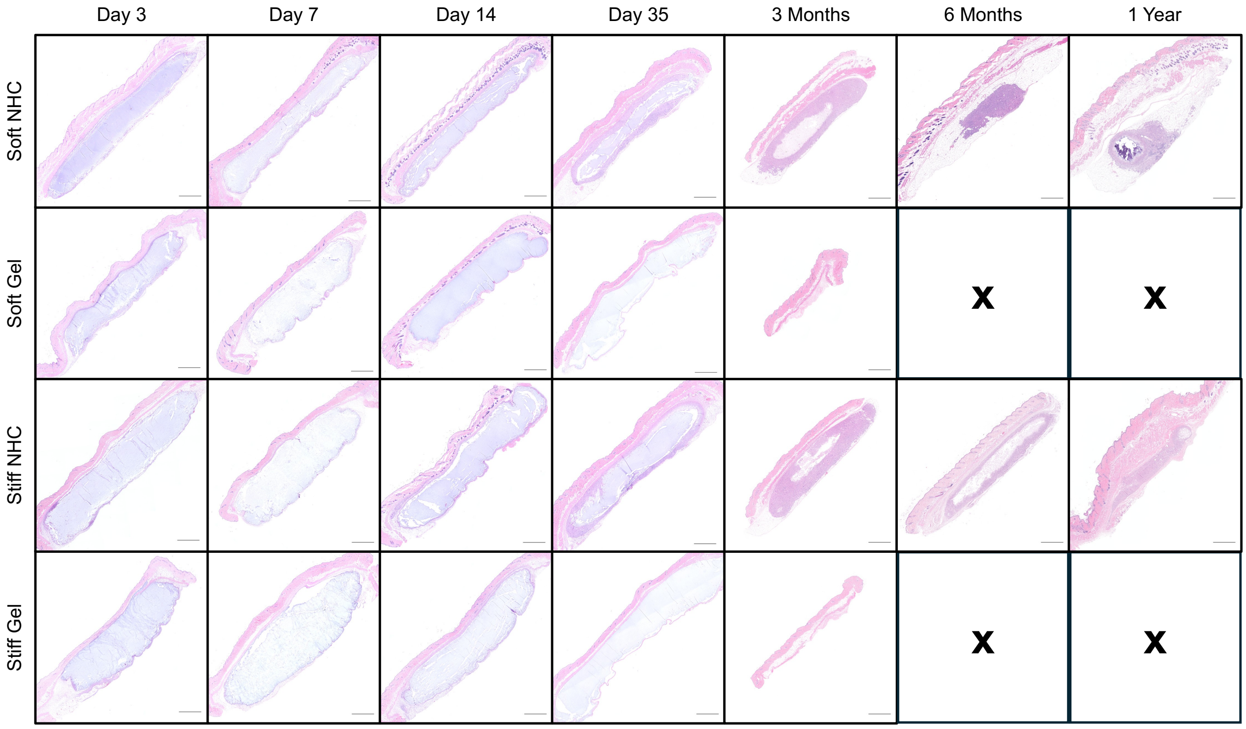


**Supplementary Figure 5. Extended representative histology.** Representative H&E-stained histologic cross-sections of materials subcutaneously injected into C57BL/6 mice, showing gross morphology including skin. Soft materials for day 3, day 7, day 14, and day 35 also appear in Figure 2a. Scale bars are 1000 μm. These H&E histological images correspond to the Masson's Trichrome-stained samples in Supplementary Figure 2.

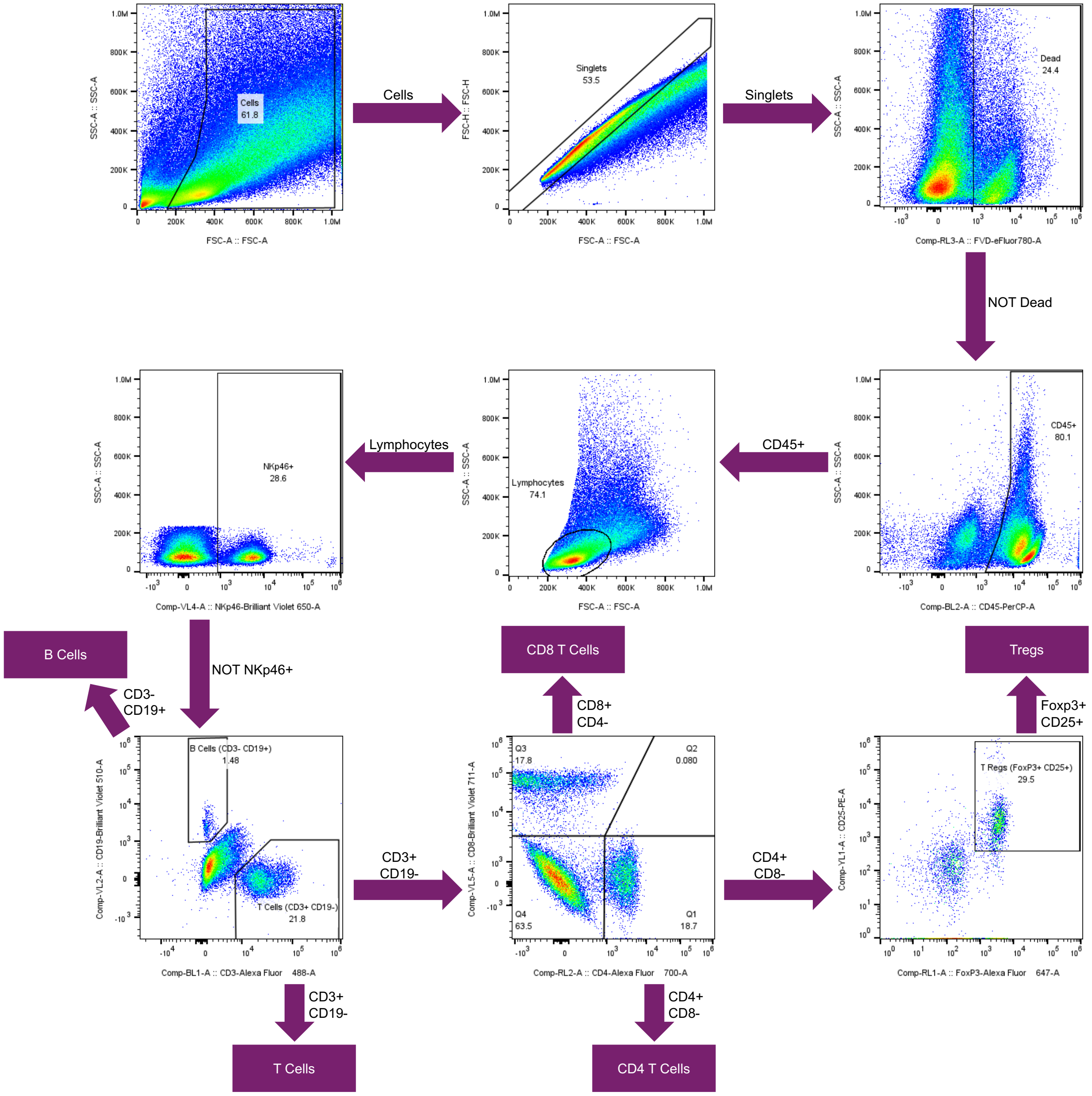


**Supplementary Figure 6. Flow cytometry gating for lymphocyte panel**. Gating scheme for flow cytometry panel designed to quantify lymphocytes.

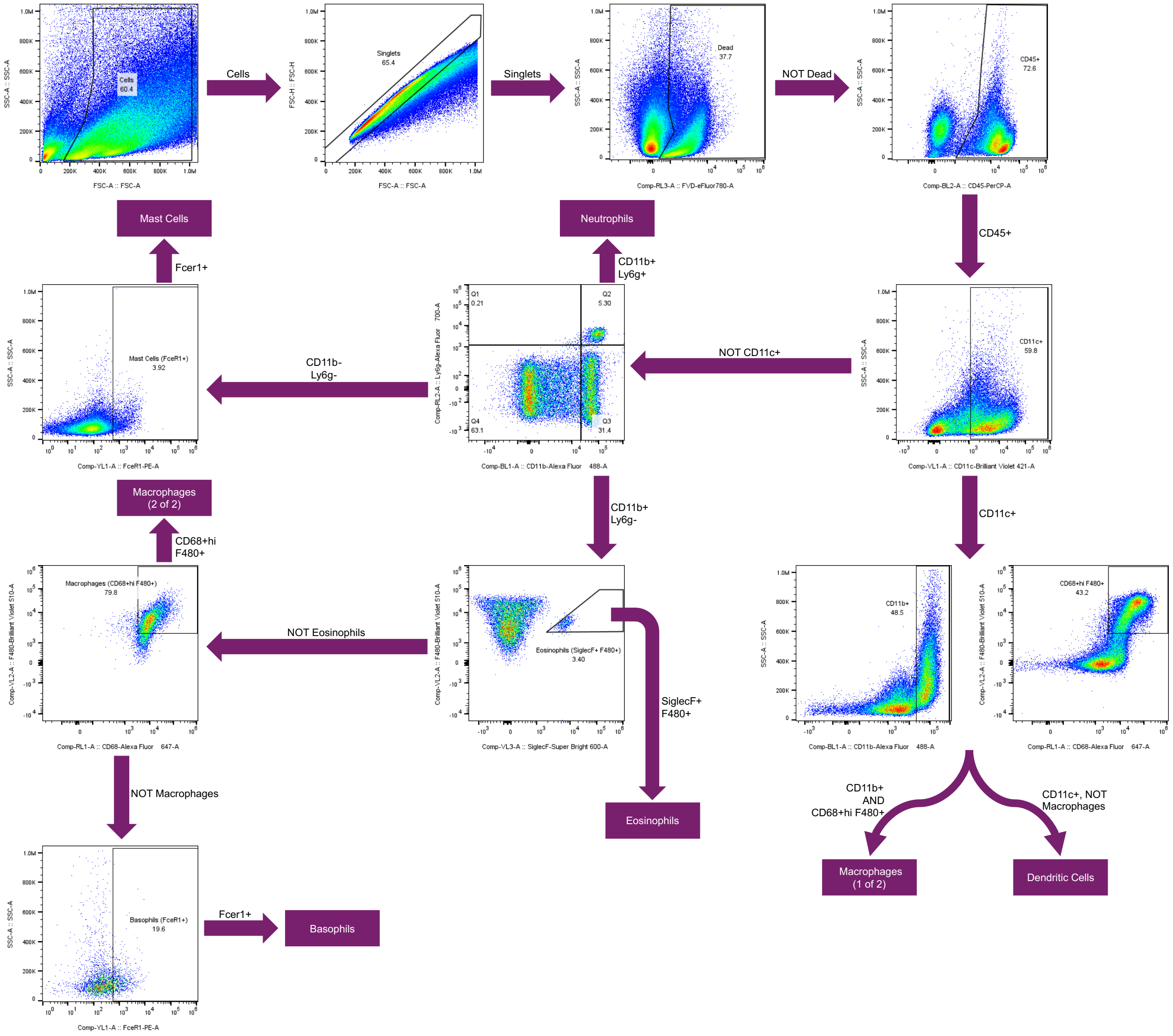


**Supplementary Figure 7. Flow cytometry gating for myeloid panel.** Gating scheme for flow cytometry panel designed to quantify myeloid immune cells.

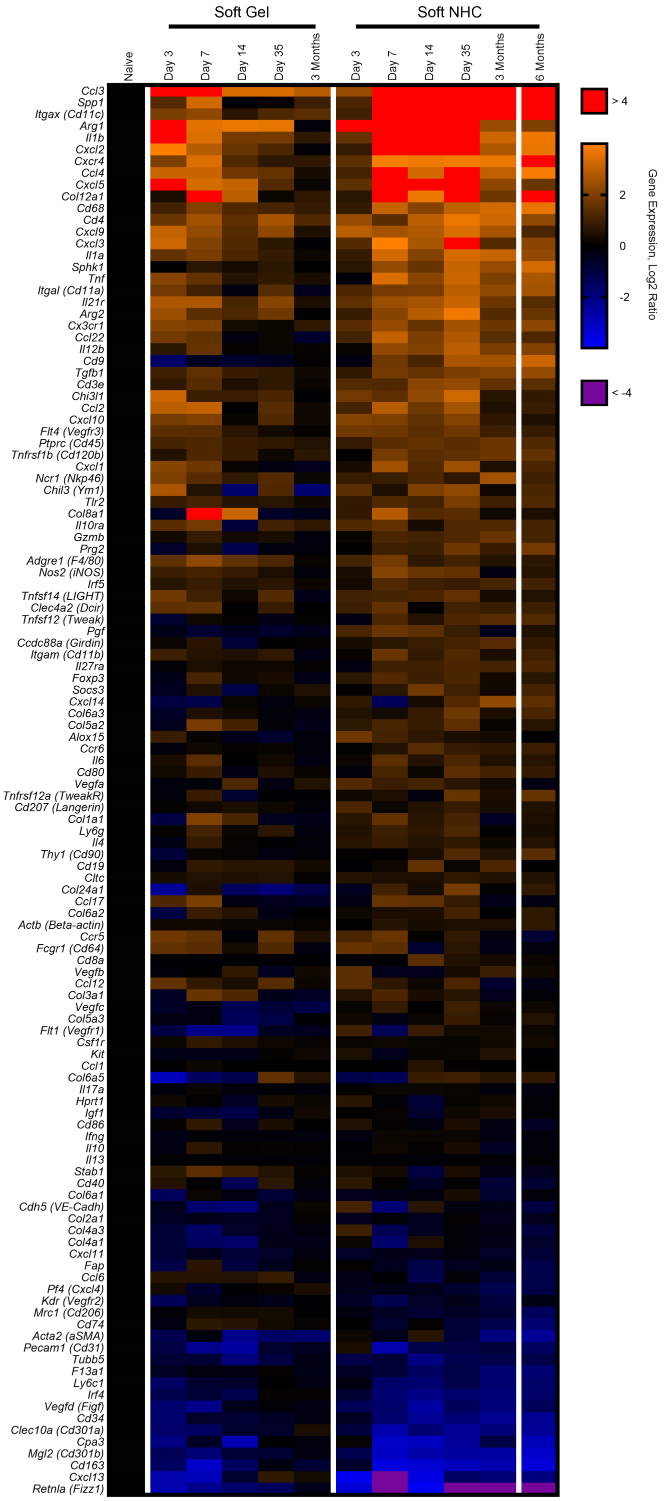


**Supplementary Figure 8. Full NanoString heatmap of all genes for WT mice.** Full heatmap showing NanoString gene expression of all genes tested for soft (150 Pa) materials explanted from C57BL/6 mice at various timepoints. The heatmap shows absolute transcript counts normalized to naïve fascia (scale: log2 ratio). Data, mean (n = 3-5 biological replicates). Genes were ordered based on average expression across all NHC groups.

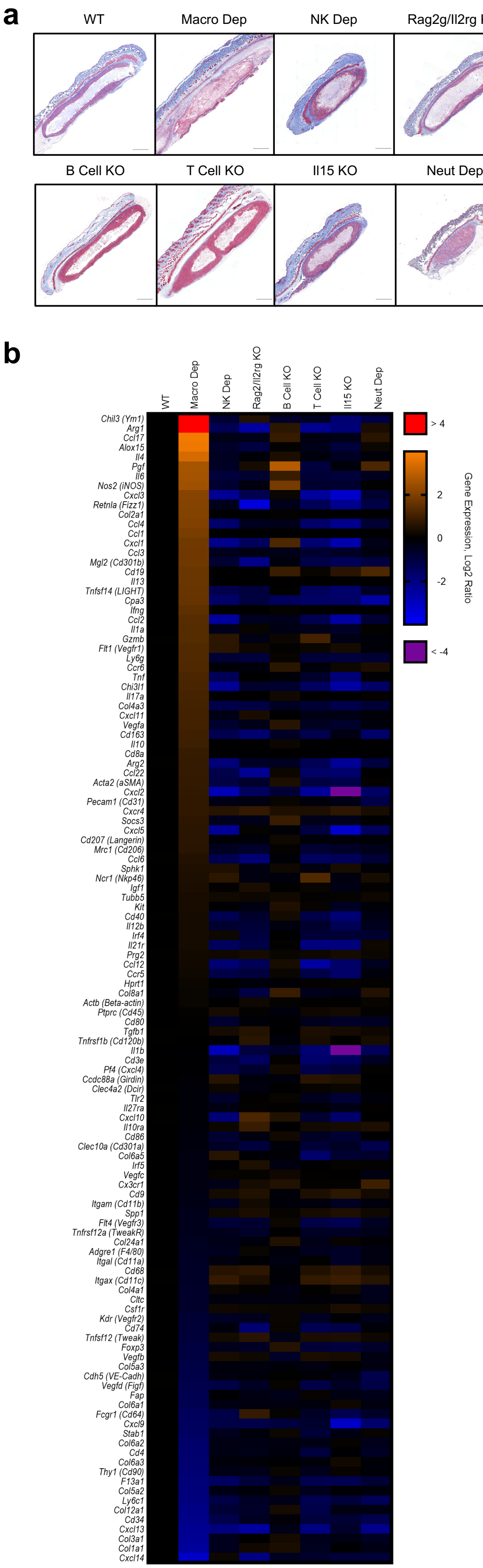
a
WT
Macro Dep
NK Dep
Rag2g/Il2rg KO
B Cell KO
T Cell KO
Il15 KO
Neut Dep
b
WT
Macro Dep
NK Dep
Rag2/Il2rg KO
B Cell KO
T Cell KO
Il15 KO
Neut Dep
Chil3 (Ym1)
Arg1
Ccl17
Alox15
Il4
Pgf
Il6
Nos2 (iNOS)
Cxcl3
Retnla (Fizz1)
Col2a1
Ccl4
Ccl1
Cxcl1
Ccl3
Mgl2 (Cd301b)
Cd19
Il13
Tnfsf14 (LIGHT)
Cpa3
Ifng
Ccl2
Il1a
Gzmb
Flt1 (Vegfr1)
Ly6g
Ccr6
Tnf
Chi3l1
Il17a
Col4a3
Cxcl11
Vegfa
Cd163
Il10
Cd8a
Arg2
Ccl22
Acta2 (aSMA)
Cxcl2
Pecam1 (Cd31)
Cxcr4
Socs3
Cxcl5
Cd207 (Langerin)
Mrc1 (Cd206)
Ccl6
Sphk1
Ncr1 (Nkp46)
Igf1
Tubb5
Kit
Cd40
Il12b
Irf4
Il21r
Prg2
Ccl12
Ccr5
Hprt1
Col8a1
Actb (Beta-actin)
Ptprc (Cd45)
Cd80
Tgfb1
Tnfrsf1b (Cd120b)
Il1b
Cd3e
Pf4 (Cxcl4)
Ccdc88a (Girdin)
Clec4a2 (Dcir)
Tlr2
Il27ra
Cxcl10
Il10ra
Cd86
Clec10a (Cd301a)
Col6a5
Irf5
Vegfc
Cx3cr1
Cd9
Itgam (Cd11b)
Spp1
Flt4 (Vegfr3)
Tnfrsf12a (TweakR)
Col24a1
Adgre1 (F4/80)
Itgal (Cd11a)
Cd68
Itgax (Cd11c)
Col4a1
Cltc
Csf1r
Kdr (Vegfr2)
Cd74
Tnfsf12 (Tweak)
Foxp3
Vegfb
Col5a3
Cdh5 (VE-Cadh)
Vegfd (Figf)
Fap
Col6a1
Fcgr1 (Cd64)
Cxcl9
Stab1
Col6a2
Cd4
Col6a3
Thy1 (Cd90)
F13a1
Col5a2
Ly6c1
Col12a1
Cd34
Cxcl13
Col3a1
Col1a1
Cxcl14
> 4
2
0
-2
< -4
Gene Expression, Log2 Ratio

**Supplementary Figure 9. Additional knockout/depletion data. a,** Representative Masson's Trichrome-stained histologic cross-sections of soft (150 Pa) NHC materials subcutaneously injected into knockout (KO)/depletion (Dep) mouse models with a C57BL/6 background, in comparison to a fully immune-competent wild type (WT) control. Materials were explanted at day 35. Mouse models shown are WT C57BL/6 mice, macrophage depletion (Macro Dep) via Clodrosome injection into C57BL/6 mice, natural killer cell depletion (NK Dep) via anti-asialo GM1 antibody injection into C57BL/6 mice, Rag2/Il2rg double knockout mice (Rag2g/Il2rg KO), muMt$^{-}$ mice (B Cell KO), B6 nude mice (T Cell KO), Il15 KO mice, and neutrophil depletion (Neut Dep) via anti-Ly6g antibody injection into C57BL/6 mice. WT, Macro Dep, and Neut Dep models also appear in Figure 3a. Scale bars are 1000 μm. **b,** Full NanoString heatmap showing gene expression of all genes tested for soft (150 Pa) materials explanted from WT or knockout/depletion C57BL/6 mice at day 35. The heatmap shows absolute transcript counts normalized to naïve fascia (scale: log2 ratio). Data, mean (n = 3-5 biological replicates). Genes were ordered based on expression in the Macro Dep group.

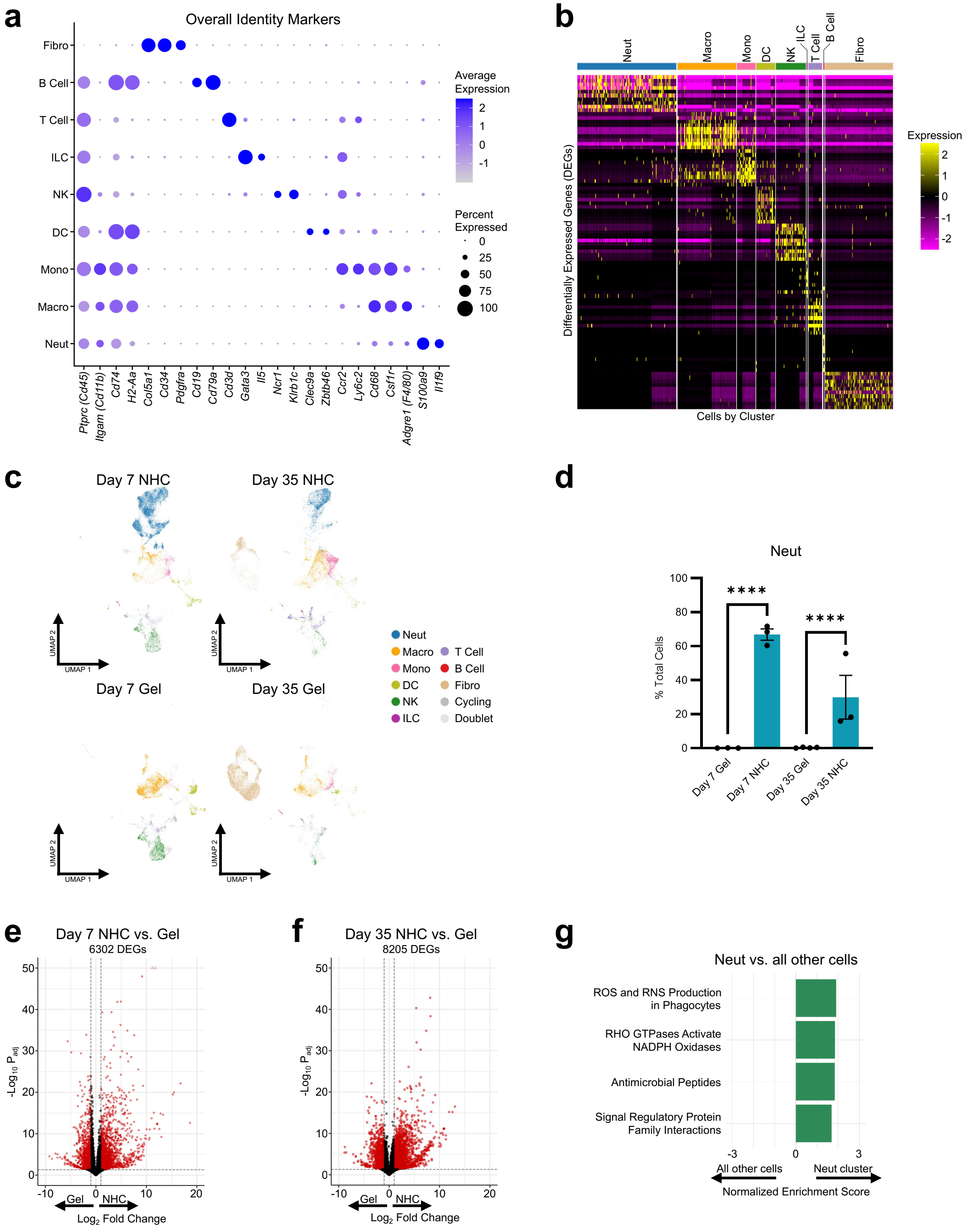


**Supplementary Figure 10. Additional scRNA-seq identity markers and neutrophil information. a,** Dot plot depicting normalized expression of canonical markers used to identify clusters. Dot size corresponds to percentage of cells within a class, and color represents the average expression level across the cells within a class. **b,** Heatmap of significant differentially expressed genes (DEGs) with the highest log fold change per cluster. **c,** UMAP plots of annotated cell clusters recruited to materials split by condition (timepoint + material). **d,** Neutrophil (Neut) differential abundance results comparing cluster abundance between groups, found using edgeR's generalized linear model framework. Data, mean ± SEM (n = 3 or 4 biological replicates/group). **e, f,** Volcano plots showing differential gene expression results, highlighting the number of significant DEGs. Pseudobulk comparisons are between NHC and gel control at day 7 (**e**) and day 35 (**f**). Red symbols represent significant genes with an adjusted p-value < 0.05 and a fold change > 1. Open triangle symbols represent values that are off scale in the direction they point. **g,** Normalized enrichment scores of select significantly enriched gene sets (adjusted *p*-value < 0.05) from Hallmark, BioCarta, WikiPathways, and Reactome collections. Neutrophil (Neut) vs. all other cells comparison was based on the Wilcoxon rank-sum test differential gene expression. In this figure's graphs, statistics shown are (*) timepoint and stiffness-matched comparisons of NHC vs. gel control. *$p < 0.05$, **$p < 0.01$, ***$p < 0.001$, ****$p < 0.0001$.

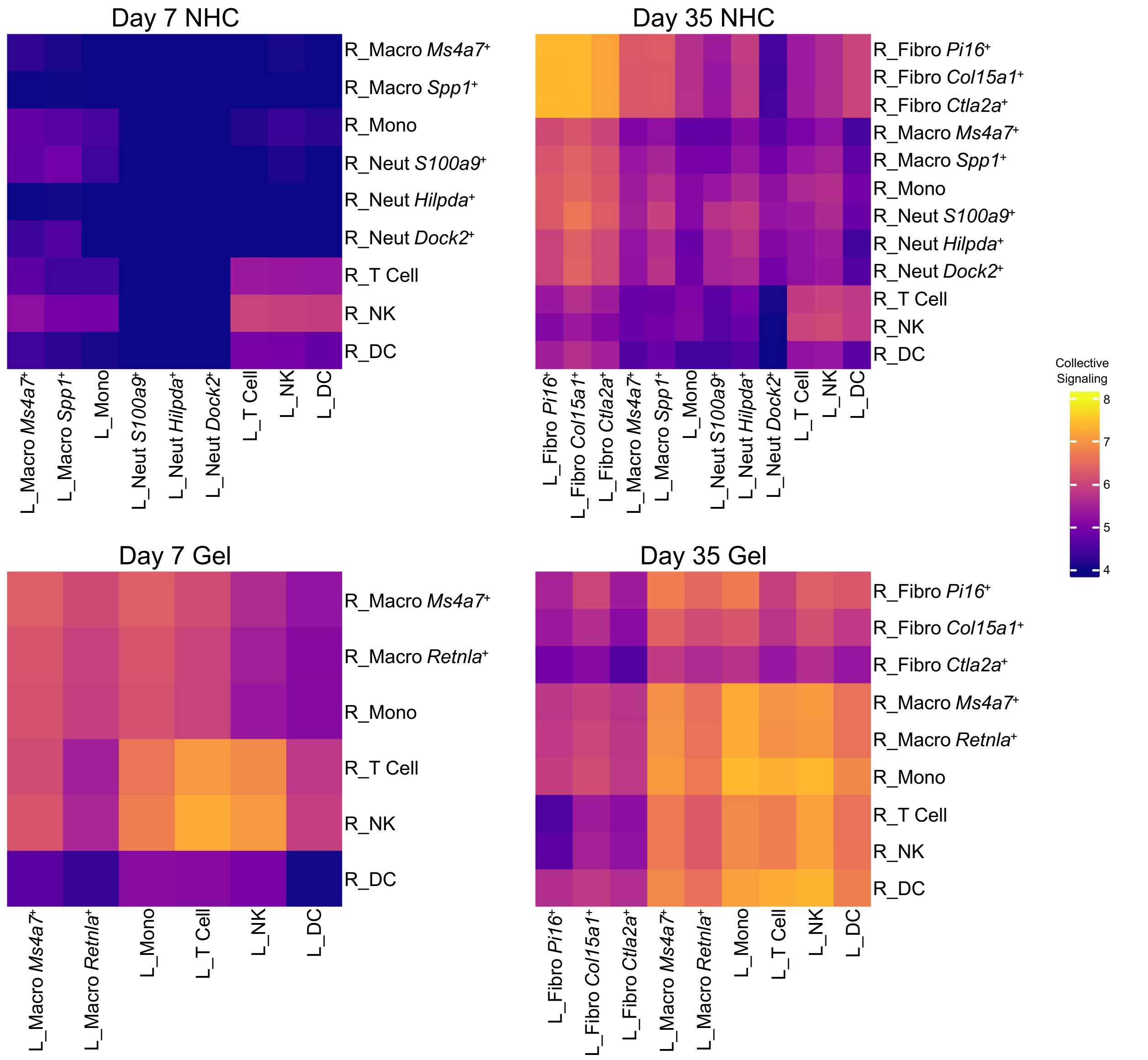


**Supplementary Figure 11. Overall communication networks showing selected subclusters of interest.** Heatmaps of inferred outgoing signaling from sending clusters (denoted "L_" for ligands) to receiving clusters (denoted "R_" for receptors) for all conditions. Color is based on the collective (summed) ligand signaling score of the sending cluster.

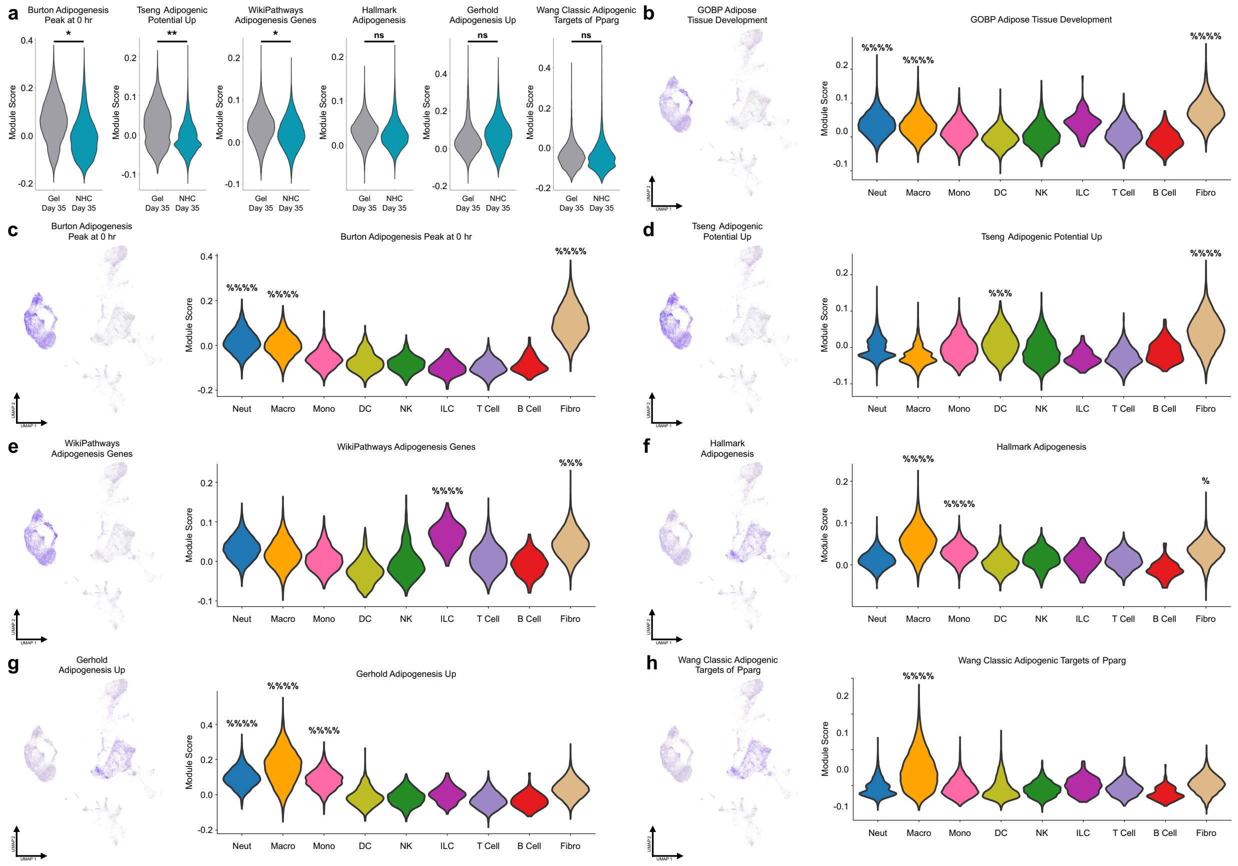


**Supplementary Figure 12. Additional adipogenesis pathways. a,** Violin plots of module score by material type at day 35 for the following pathways: "Burton Adipogenesis Peak at 0hr", "Tseng Adipogenesis Potential Up", "WP Adipogenesis Genes", "Hallmark Adipogenesis", "Gerhold Adipogenesis Up", and "Wang Classic Adipogenic Targets of Pparg". Independent t-test. **b–h,** feature plot showing module score of genes most correlated to pathway (*left*) and violin plot by cluster showing module score (*right*) for both NHC and gel control samples at day 35. Plots are for the following pathways: "GOBP Adipose Tissue Development" (**b**), "Burton Adipogenesis Peak at 0hr" (**c**), "Tseng Adipogenesis Potential Up" (**d**), "WP Adipogenesis Genes" (**e**), "Hallmark Adipogenesis" (**f**), "Gerhold Adipogenesis Up" (**g**), and "Wang Classic Adipogenic Targets of Pparg" (**h**). Independent t-test with Benjamini-Hochberg correction for multiple comparisons. Significance only shown for clusters with higher expression than the average expression of all other clusters. In this figure's graphs, statistics shown are (*) timepoint and stiffness-matched comparisons of NHC vs. gel control, as well as ($^{\%}$) comparisons of a single cluster vs. all other cells. *,$^{\%}$:$p < 0.05$; **,$^{\%\%}$:$p < 0.01$; ***,$^{\%\%\%}$:$p < 0.001$; ****,$^{\%\%\%\%}$:$p < 0.0001$.

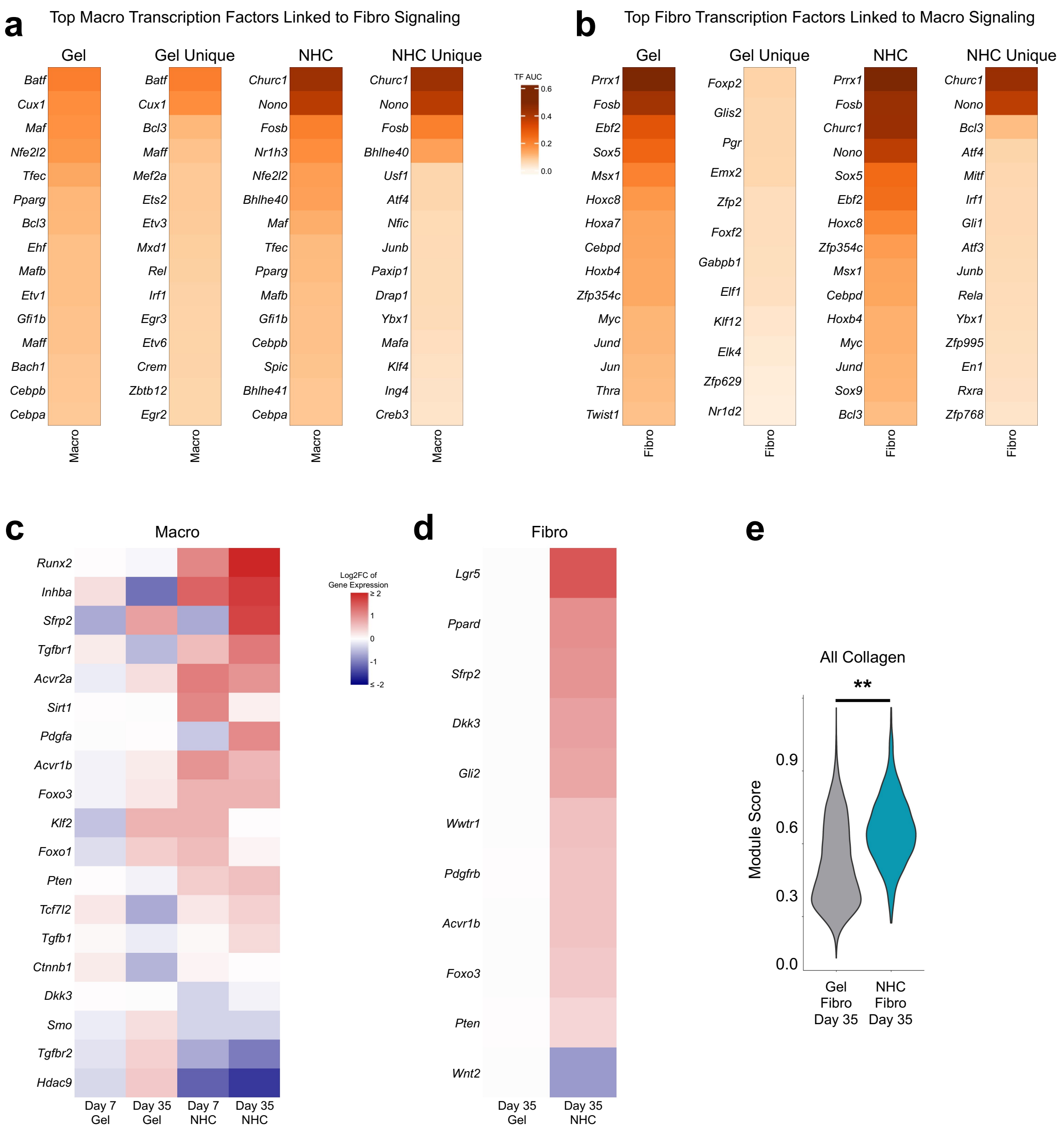


**Supplementary Figure 13. NHC materials influence early adipogenesis-related signaling during niche establishment. a, b,** Heatmap of top cluster-averaged pySCENIC-calculated transcription factor area under the curve (TF AUC) scores. Transcription factors shown were found by DominoSignal to be activated in macrophages by fibroblasts (**a**) or activated in fibroblasts by macrophages (**b**). From left to right, heatmaps show the top TF AUCs in gel control, the top unique TF AUCs in gel control, the top TF AUCs in NHC, and the top unique TF AUCs in NHC. **c, d,** Heatmap of cluster-averaged fold change in gene expression vs. gel control for select significant (adjusted $p$-value < 0.05) differentially expressed genes between NHC and gel control at timepoint-matched comparisons (scale: log2 fold change). Genes shown are within selected pySCENIC transcription factor gene sets that were implicated by DominoSignal in macrophage-fibroblast signaling and related to negative regulation of adipogenesis. Genes were ordered based on average expression in NHC. Heatmaps are shown for macrophages (**c**) and fibroblasts (**d**). **e**, Violin plot of module score by material type at day 35 for fibroblasts for all collagen genes (*Col1a1*,

*Col1a2*, *Col2a1*, *Col3a1*, *Col4a1*, *Col4a2*, *Col4a3*, *Col4a4*, *Col4a5*, *Col4a6*, *Col5a1*, *Col5a2*, *Col5a3*, *Col6a1*, *Col6a2*, *Col6a3*, *Col6a4*, *Col6a5*, *Col6a6*, *Col7a1*, *Col8a1*, *Col8a2*, *Col9a1*, *Col9a2*, *Col9a3*, *Col10a1*, *Col11a1*, *Col11a2*, *Col12a1*, *Col13a1*, *Col14a1*, *Col15a1*, *Col16a1*, *Col17a1*, *Col18a1*, *Col19a1*, *Col20a1*, *Col22a1*, *Col23a1*, *Col24a1*, *Col25a1*, *Col26a1*, *Col27a1*, & *Col28a1*). Independent t-test. In this figure's graphs, statistics shown are (*) timepoint and stiffness-matched comparisons of NHC vs. gel control. *$p < 0.05$, **$p < 0.01$, ***$p < 0.001$, ****$p < 0.0001$.

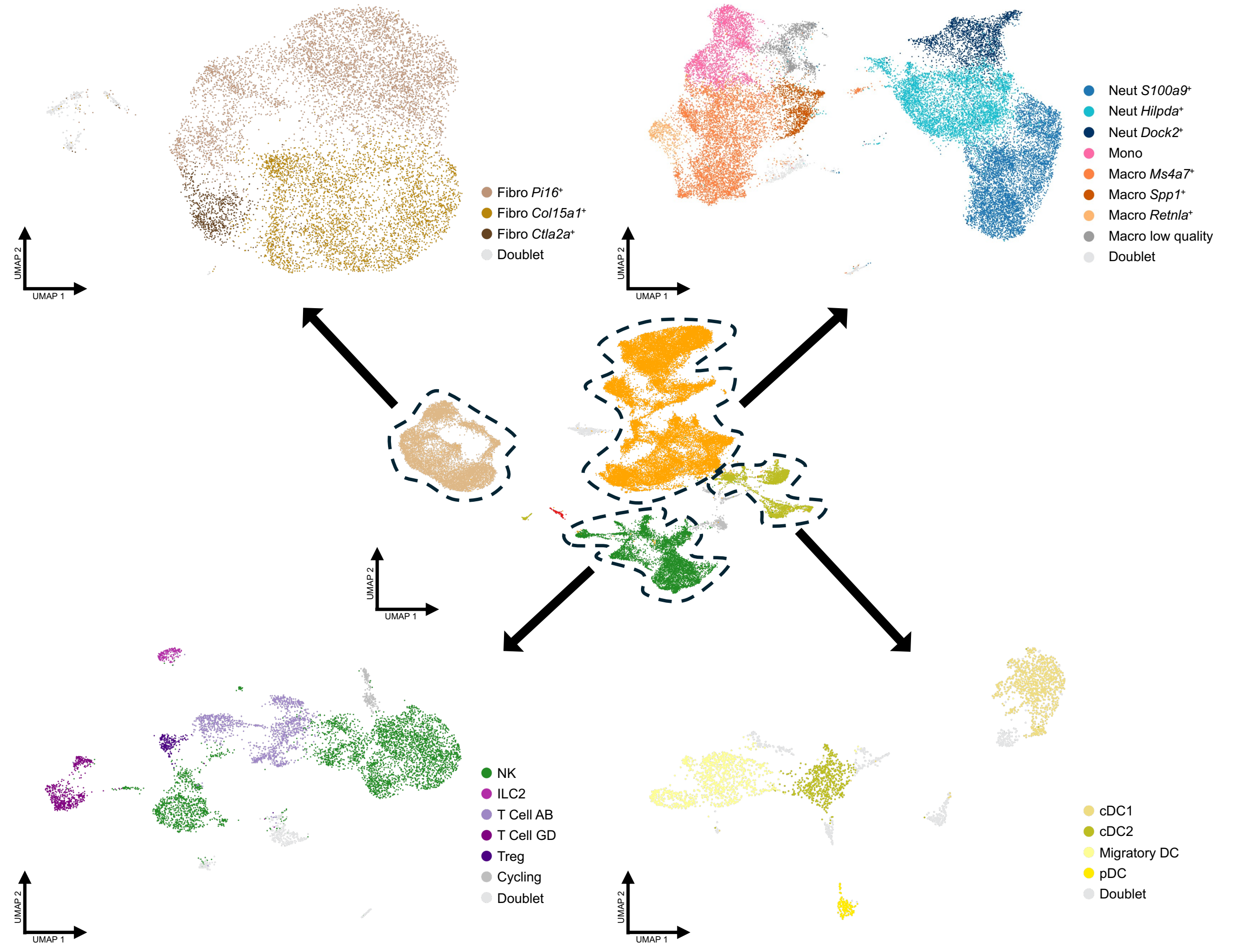


**Supplementary Figure 14. Clustering and subclustering of scRNA-seq data.** UMAP plots of the initial clustering (*middle*) and subsequent subclustering (*outer corners*) performed on the scRNA-seq dataset.

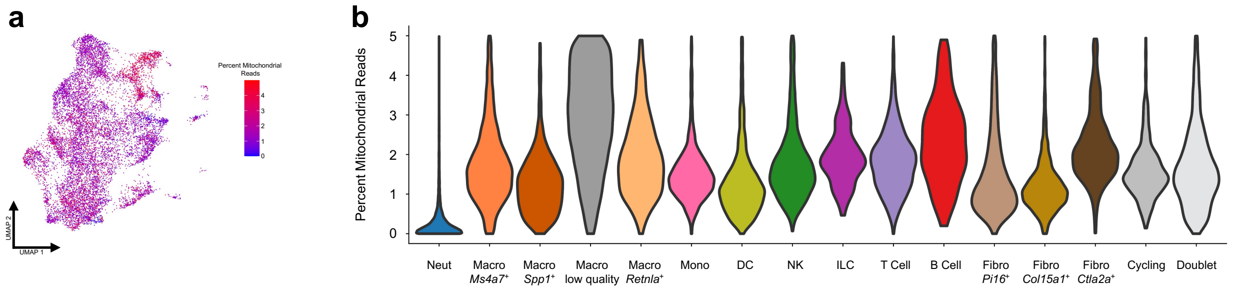


**Supplementary Figure 15. Exclusion of low-quality macrophages. a,** Feature plot showing percent mitochondrial reads in the macrophages and monocytes. **b,** Violin plot of percent mitochondrial reads by cluster/subcluster.

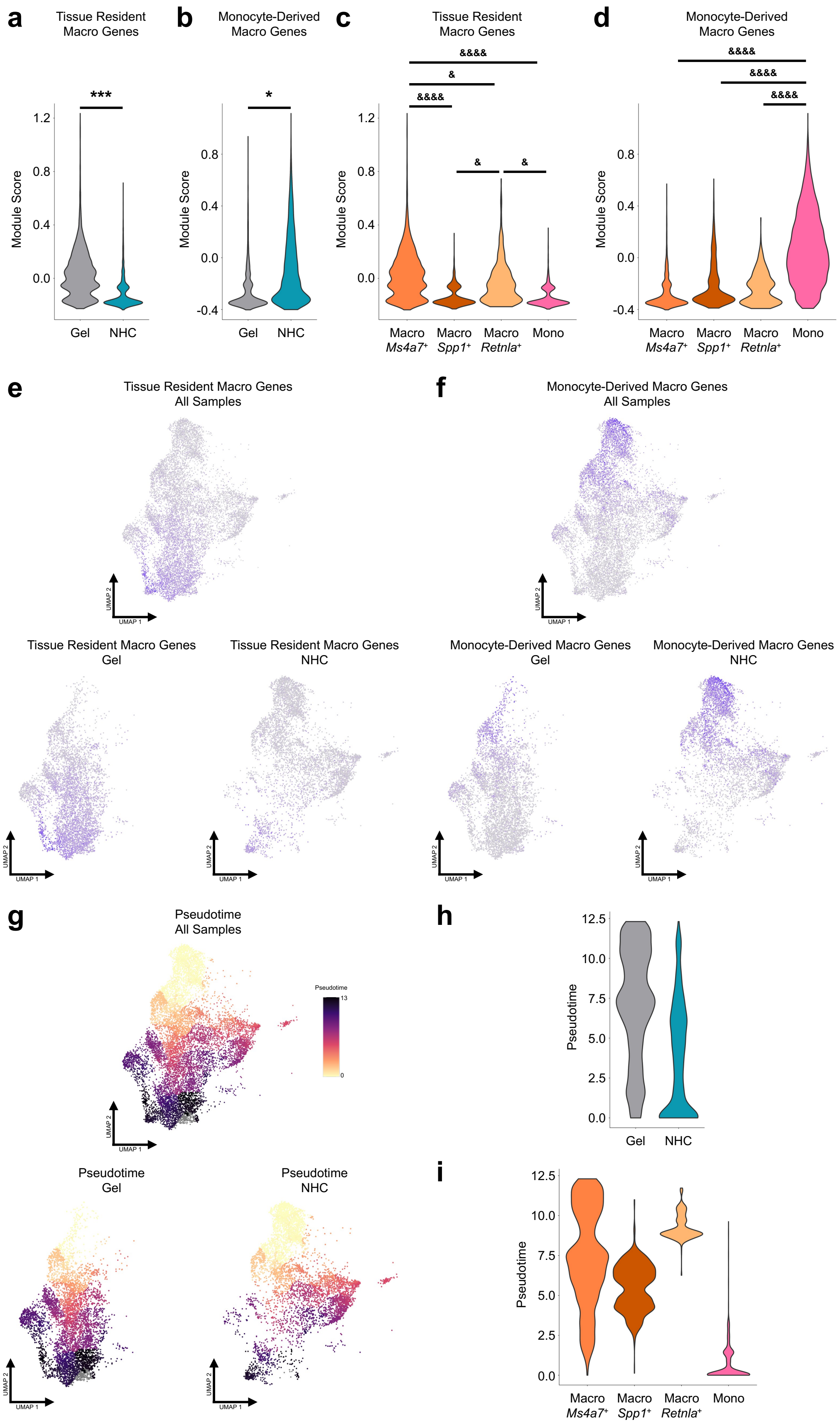

a
Tissue Resident Macro Genes
b
Monocyte-Derived Macro Genes
c
Tissue Resident Macro Genes
d
Monocyte-Derived Macro Genes
Module Score
Gel
NHC
Macro Ms4a7+
Macro Spp1+
Macro Retnla+
Mono
e
Tissue Resident Macro Genes All Samples
f
Monocyte-Derived Macro Genes All Samples
Tissue Resident Macro Genes Gel
Tissue Resident Macro Genes NHC
Monocyte-Derived Macro Genes Gel
Monocyte-Derived Macro Genes NHC
g
Pseudotime All Samples
Pseudotime Gel
Pseudotime NHC
h
i
Pseudotime

**Supplementary Figure 16. Macrophage trajectory inference reveals skewed populations by material type. a, b,** Violin plots of module scores by material type for tissue resident macrophage genes (**a**) and monocyte-derived macrophage genes (**b**). Independent t-test. **c, d,** Violin plots of module scores by relevant subcluster for tissue resident macrophage genes (**c**) and monocyte-derived macrophage genes (**d**). One-way ANOVA followed by Tukey's multiple comparisons test. **e, f,** Feature plots showing module scores for tissue resident macrophage genes (**e**) and monocyte-derived macrophage genes (**f**) in all samples (*top middle*), gel control (*bottom left*), and NHC (*bottom right*). **g,** Monocle3-inferred pseudotime trajectory of macrophages and monocytes, showing the spectrum from monocytes to monocyte-derived-like macrophages to tissue resident-like macrophages in all samples (*top middle*), gel control (*bottom left*), and NHC (*bottom right*). **h,** Violin plot of pseudotime values for the monocyte to tissue-resident-like macrophage trajectory by material type. **i,** Violin plot of pseudotime values for the monocyte to tissue-resident-like macrophage trajectory by relevant subcluster. In this figure, the tissue-resident macrophage marker genes used were *Adgre1*, *Timd4*, *Folr2*, *Lyve1*, *Cd163*, & *Vsig4*; the monocyte-derived macrophage marker genes used were *Myb*, *Ms4a3*, *Ccr2*, *Ly6c2*, & *Sell*. In this figure's graphs, statistics shown are (*) timepoint and stiffness-matched comparisons of NHC vs. gel control, as well as ($^{\&}$) comparisons between clusters. *,$^{\&}$:$p < 0.05$; **,$^{\&\&}$:$p < 0.01$; ***,$^{\&\&\&}$:$p < 0.001$; ****,$^{\&\&\&\&}$:$p < 0.0001$.

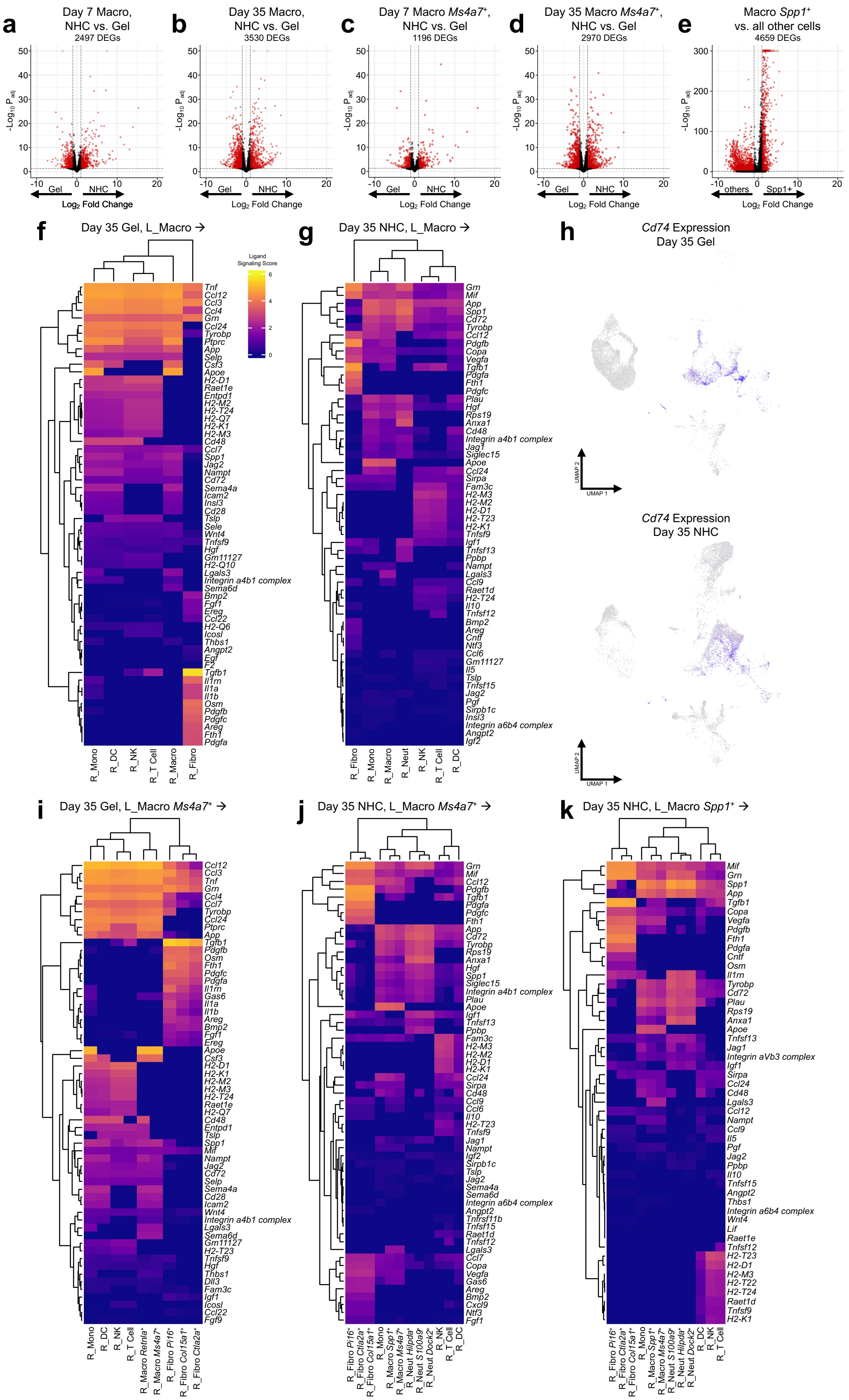
a
Day 7 Macro,
NHC vs. Gel
2497 DEGs
b
Day 35 Macro,
NHC vs. Gel
3530 DEGs
c
Day 7 Macro *Ms4a7*⁺,
NHC vs. Gel
1196 DEGs
d
Day 35 Macro *Ms4a7*⁺,
NHC vs. Gel
2970 DEGs
e
Macro *Spp1*⁺
vs. all other cells
4659 DEGs
$-\text{Log}_{10}\ P_{adj}$
Gel
NHC
others
Spp1+
$\text{Log}_2$ Fold Change
f
Day 35 Gel, L_Macro →
Ligand Signaling Score
g
Day 35 NHC, L_Macro →
h
*Cd74* Expression
Day 35 Gel
*Cd74* Expression
Day 35 NHC
UMAP 1
UMAP 2
R_Mono
R_DC
R_NK
R_T Cell
R_Macro
R_Fibro
R_Neut
i
Day 35 Gel, L_Macro *Ms4a7*⁺ →
j
Day 35 NHC, L_Macro *Ms4a7*⁺ →
k
Day 35 NHC, L_Macro *Spp1*⁺ →
R_Macro *Retnla*⁺
R_Macro *Ms4a7*⁺
R_Fibro *Pi16*⁺
R_Fibro *Col15a1*⁺
R_Fibro *Ctla2a*⁺
R_Macro *Spp1*⁺
R_Neut *Hilpda*⁺
R_Neut *S100a9*⁺
R_Neut *Dock2*⁺

**Supplementary Figure 17. Extended scRNA-seq analysis for macrophages. a–e,** Volcano plots showing differential gene expression results, highlighting the number of significant differentially expressed genes (DEGs). Pseudobulk comparisons are between NHC and gel control for day 7 macrophages (**a**), day 35 macrophages (**b**), day 7 *Ms4a7*$^+$ macrophages (**c**), and day 35 *Ms4a7*$^+$ macrophages (**d**). Wilcoxon rank-sum test comparison between a cluster and all other cells is for *Spp1*$^+$ macrophages (**e**). Red symbols represent significant genes with an adjusted *p*-value < 0.05 and a fold change > 1. Open triangle symbols represent values that are off scale in the direction they point. **f, g,** Heatmaps of inferred outgoing signaling for all ligands at day 35 from the macrophage cluster in gel control (**f**) and NHC (**g**). Sending cluster(s) are denoted "L" for ligands and receiving cluster(s) are denoted "R_" for receptors. Color is based on ligand signaling scores. **h,** Feature plots showing normalized gene expression for receptor *Cd74* in day 35 gel control (*top*) and NHC (*bottom*). **i–k,** Heatmaps of inferred outgoing signaling for all ligands at day 35 from the *Ms4a7*$^+$ macrophage subcluster in gel control (**i**), *Ms4a7*$^+$ macrophage subcluster in NHC (**j**), and *Spp1*$^+$ macrophage subcluster in NHC (**k**). Sending cluster(s) are denoted "L" for ligands and receiving cluster(s) are denoted "R_" for receptors. Color is based on ligand signaling scores.

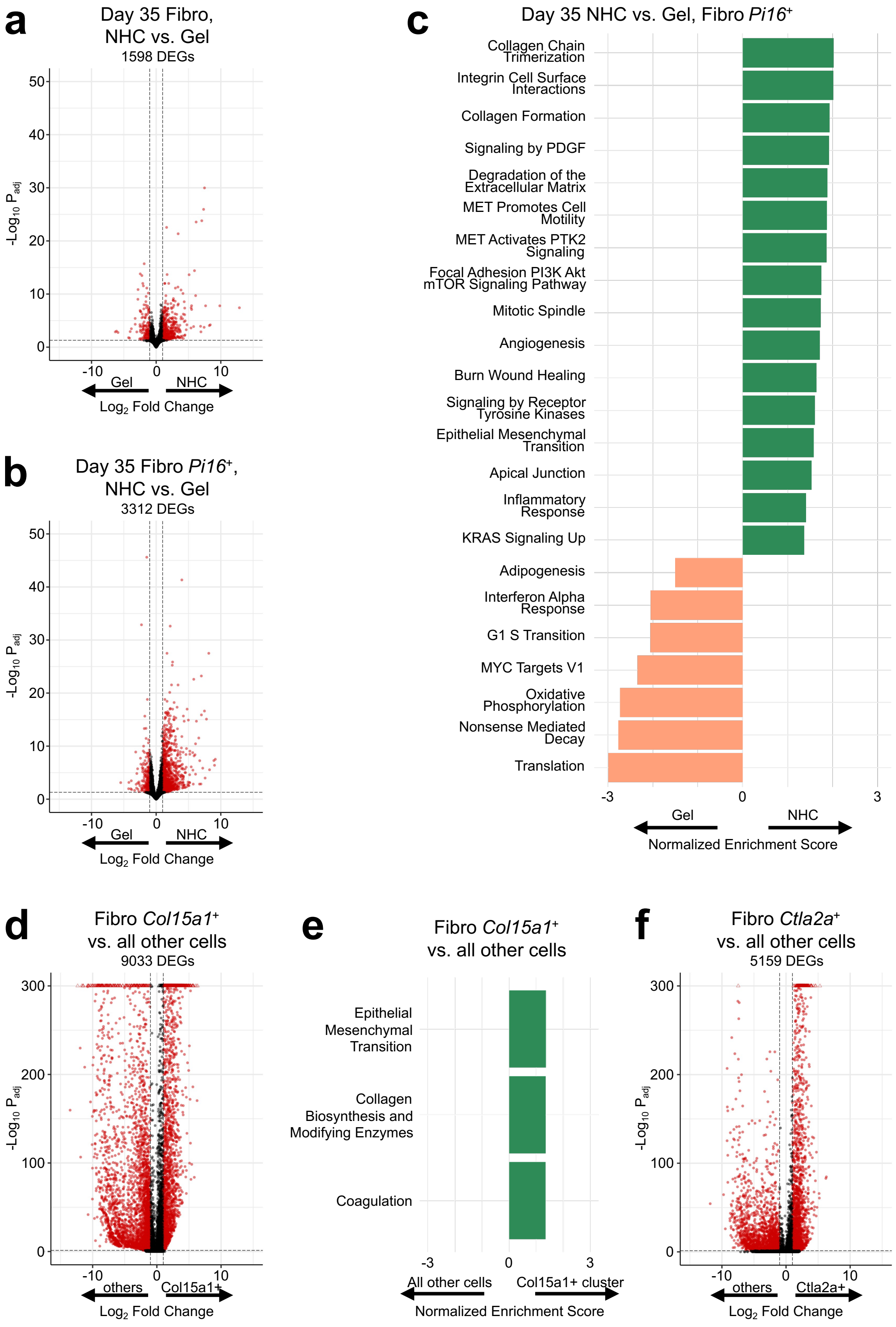
a
Day 35 Fibro, NHC vs. Gel
1598 DEGs
-Log10 Padj
Gel
NHC
Log2 Fold Change
b
Day 35 Fibro Pi16+, NHC vs. Gel
3312 DEGs
c
Day 35 NHC vs. Gel, Fibro Pi16+
Collagen Chain Trimerization
Integrin Cell Surface Interactions
Collagen Formation
Signaling by PDGF
Degradation of the Extracellular Matrix
MET Promotes Cell Motility
MET Activates PTK2 Signaling
Focal Adhesion PI3K Akt mTOR Signaling Pathway
Mitotic Spindle
Angiogenesis
Burn Wound Healing
Signaling by Receptor Tyrosine Kinases
Epithelial Mesenchymal Transition
Apical Junction
Inflammatory Response
KRAS Signaling Up
Adipogenesis
Interferon Alpha Response
G1 S Transition
MYC Targets V1
Oxidative Phosphorylation
Nonsense Mediated Decay
Translation
Normalized Enrichment Score
d
Fibro Col15a1+ vs. all other cells
9033 DEGs
others
Col15a1+
e
Fibro Col15a1+ vs. all other cells
Epithelial Mesenchymal Transition
Collagen Biosynthesis and Modifying Enzymes
Coagulation
All other cells
Col15a1+ cluster
f
Fibro Ctla2a+ vs. all other cells
5159 DEGs
Ctla2a+

**Supplementary Figure 18. Extended scRNA-seq differential gene expression and gene set enrichment analysis for fibroblasts. a, b,** Volcano plots showing differential gene expression results, highlighting the number of significant differentially expressed genes (DEGs). Pseudobulk comparisons are between NHC and gel control for day 35 fibroblasts (**a**) and day 35 *Pi16*$^{+}$ fibroblasts (**b**). Red symbols represent significant genes with an adjusted *p*-value < 0.05 and a fold change > 1. Open triangle symbols represent values that are off scale in the direction they point. **c,** Normalized enrichment scores of select significantly enriched gene sets (adjusted *p*-value < 0.05) from Hallmark, BioCarta, WikiPathways, and Reactome collections. NHC vs. gel control comparison based on pseudobulk differential gene expression is for day 35 *Pi16*$^{+}$ fibroblasts. **d,** Volcano plot showing differential gene expression results, highlighting the number of significant DEGs. Wilcoxon rank-sum test comparison between *Col15a1*$^{+}$ fibroblasts and all other cells. Red symbols represent significant genes with an adjusted *p*-value < 0.05 and a fold change > 1. Open triangle symbols represent values that are off scale in the direction they point. **e,** Normalized enrichment scores of select significantly enriched gene sets (adjusted *p*-value < 0.05) from Hallmark, BioCarta, WikiPathways, and Reactome collections. Cluster vs. all other cells comparison based on the Wilcoxon rank-sum test differential gene expression is for *Col15a1*$^{+}$ fibroblasts. **f,** Volcano plot showing differential gene expression results, highlighting the number of significant DEGs. Wilcoxon rank-sum test comparison between *Ctla2a*$^{+}$ fibroblasts and all other cells. Red symbols represent significant genes with an adjusted *p*-value < 0.05 and a fold change > 1. Open triangle symbols represent values that are off scale in the direction they point.

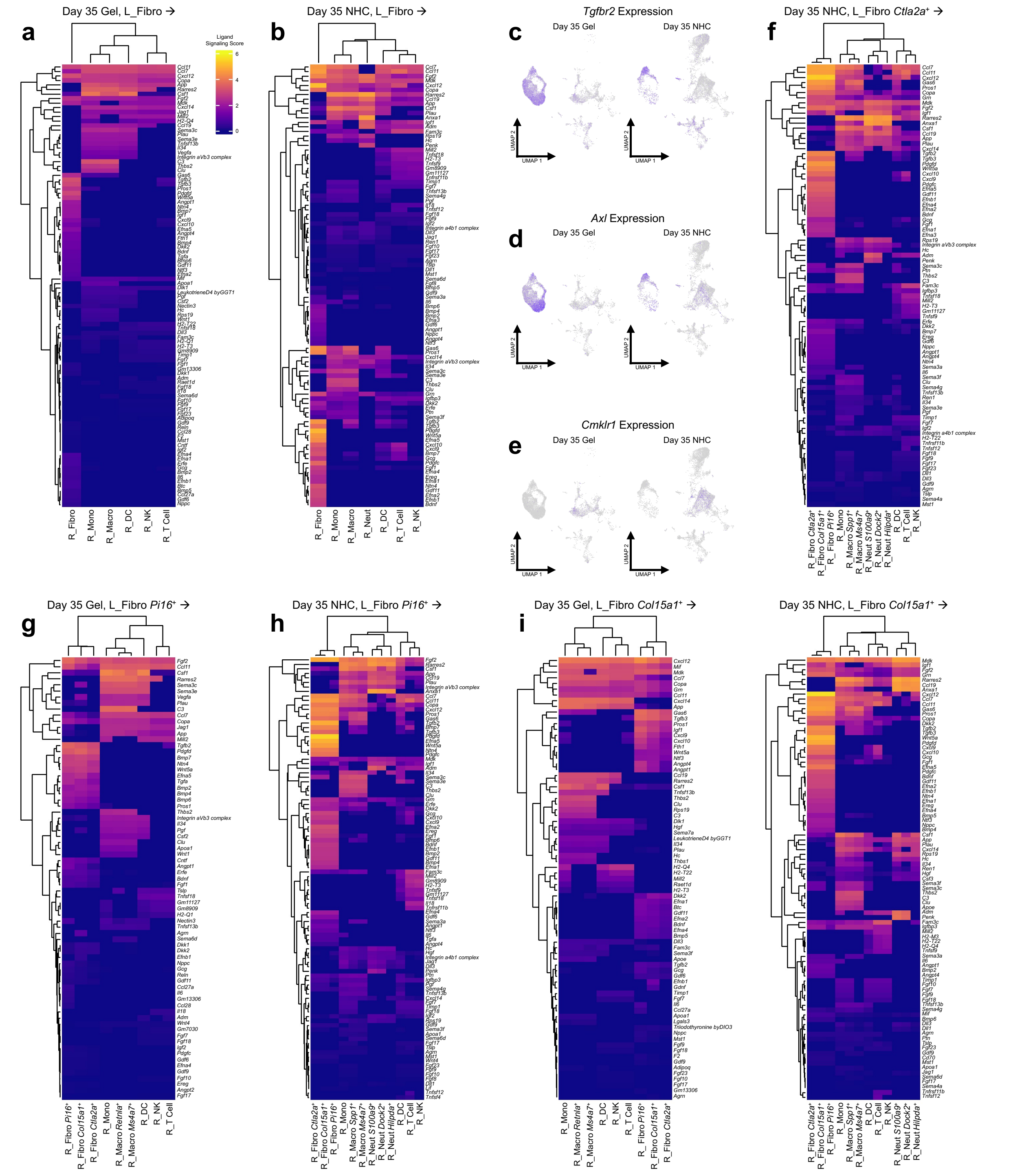


**Supplementary Figure 19. Extended scRNA-seq cell signaling analysis for fibroblasts. a, b,** Heatmaps of inferred outgoing signaling for all ligands at day 35 from the fibroblast cluster in gel control (**a**) and NHC (**b**). Sending cluster(s) are denoted “L” for ligands and receiving cluster(s) are denoted “R_” for receptors. Color is based on ligand signaling scores. **c–e,** Feature plots showing normalized gene expression for receptors *Tgfbr2*, *Axl*, and *Cmklr1* in day 35 gel control (left) and NHC (right). **f–j,** Heatmaps of inferred outgoing signaling for all ligands at day 35 from the *Ctla2a*$^{+}$ fibroblast subcluster in NHC (**f**), *Pi16*$^{+}$ fibroblast subcluster in gel control (**g**), *Pi16*$^{+}$

fibroblast subcluster in NHC (**h**), *Col15a1*$^+$ fibroblast subcluster in gel control (**i**), and *Col15a1*$^+$ fibroblast subcluster in NHC (**j**). Sending cluster(s) are denoted "L" for ligands and receiving cluster(s) are denoted "R_" for receptors. Color is based on ligand signaling scores.

### Supplementary Tables

**Supplementary Table 1. Flow Cytometry Cell Population Abundance.** Flow cytometric analysis of cells recruited to materials explanted from C57BL/6 mice at various timepoints, shown as $\%CD45^+$ cells. Data, mean (n = 5 of biological replicates). Two-way ANOVA followed by Tukey's multiple comparisons test. Immune cell types were defined as follows: dendritic cells ($CD45^+CD11c^+$), macrophages ($CD45^+CD11b^+CD68^+F4/80^+$), basophils ($CD45^+CD11b^+FceR1^+$), mast cells ($CD45^+CD11b^-FceR1^+$), neutrophils ($CD45^+CD11b^+Ly6g^+$), eosinophils ($CD45^+Cd11b^+SiglecF^+F4/80^+$), natural killer cells ($CD45^+NKp46^+$), B cells ($CD45^+CD19^+CD3^-$), & T cells ($CD45^+CD3^+CD19^-$). Statistics shown are (*) timepoint and stiffness-matched comparisons of NHC vs. gel control. *$p < 0.05$, **$p < 0.01$, ***$p < 0.001$, ****$p < 0.0001$. Flow cytometry data is graphed in Figure 2d.

| | Day 3 | | Day 7 | | Day 35 | | 3 Months | |
|---|---|---|---|---|---|---|---|---|
| | **Soft Gel** | **Soft NHC** | **Soft Gel** | **Soft NHC** | **Soft Gel** | **Soft NHC** | **Soft Gel** | **Soft NHC** |
| **Dendritic Cells** | *7.69 | *17.42 | ***23.92 | ***38.56 | ***35.88 | ***22.68 | 28.99 | 32.20 |
| **Macrophages** | 1.85 | 5.04 | 8.68 | 15.54 | 30.81 | 27.21 | 24.94 | 17.56 |
| **Basophils** | 1.00 | 1.49 | 1.01 | 2.08 | 1.37 | 2.74 | ****5.10 | ****1.84 |
| **Mast Cells** | 0.50 | 0.44 | 0.35 | 1.08 | *1.54 | *0.60 | 1.07 | 0.23 |
| **Neutrophils** | 58.73 | 55.11 | *35.15 | *10.26 | 0.57 | 5.81 | 12.55 | 4.64 |
| **Eosinophils** | 0.28 | 0.88 | 1.01 | 1.31 | 0.09 | 0.35 | 0.69 | 0.41 |
| **Natural Killer Cells** | 0.23 | 4.39 | 10.99 | 13.85 | ***19.81 | ***10.95 | 7.13 | 10.95 |
| **B Cells** | 0.01 | 0.10 | 0.13 | 0.42 | 0.80 | 1.69 | *0.30 | *1.68 |
| **T Cells** | 0.10 | 1.76 | 6.40 | 4.95 | 9.13 | 9.24 | 5.40 | 10.98 |
| **Other** | 29.60 | 13.37 | 12.36 | 11.95 | 0.00 | 18.73 | 13.83 | 19.50 |
